# Two distinct types of nodes of Ranvier support auditory nerve function in the mouse cochlea

**DOI:** 10.1101/2021.02.07.430087

**Authors:** Clarisse H. Panganiban, Jeremy L. Barth, Junying Tan, Kenyaria V. Noble, Carolyn M. McClaskey, James W. Dias, Kelly C. Harris, Hainan Lang

**Author notes:** Corresponding author:* Dr. Hainan Lang, Department of Pathology and Laboratory Medicine, Medical University of South Carolina, 165 Ashley Avenue, PO BOX 250908, Charleston, SC 29425, USA. Author contributions:* C.H.P. and H.L. designed the research; C.H.P., J.L.B., J.T., K.V.N., C.M., and H.L., performed the research; C.H.P., H.L., K.H, J.D., and J.L.B. analyzed data; and C.H.P. and H.L. wrote the paper.

## Abstract

Glial cells of the auditory nerve regulate formation of the nodes of Ranvier that are needed for regeneration of action potentials and proper hearing function. Here we identify and describe the distinct features of two novel types of Ranvier nodes—the axonal node and the ganglion node—in the mouse auditory nerve that change across the lifespan, including during myelination and postnatal development, and degenerate during aging. Cellular, molecular, and structure-function correlation evaluations revealed that nodal types are critical for different aspects of auditory nerve function. Specifically, the length of the axonal node is associated with neural processing speed and neural synchrony, whereas ganglion node development is associated with amplitude growth of the action potential. Moreover, our data indicate that dysregulation of glial cells and associated degeneration of the ganglion node structure are an important and new mechanism of auditory nerve dysfunction in age-related hearing loss.

## Introduction

Proper passage of electrical impulses and the regulation of conduction velocity through the peripheral auditory nerve (AN) are crucial for effective, precise transfer of sound information from the cochlea to the brainstem. Cochlear glial cells enwrap AN fibers and cell bodies with multi-layered myelin sheaths during early postnatal development (Romand & Romand, 1990). In addition to providing electrical insulation, the formation of myelin supports for the assembly of nodes of Ranvier at the naked gaps between myelin sheaths along axons. Node of Ranvier is made up of three separate microdomains: 1) the node, which contains clusters of voltage-gated sodium channels required for action potential regeneration, 2) the paranode flanks, where the myelin lamellae terminate and provide a physical barrier between ion channels in the node and juxtaparanodes, and 3) the juxtaparanodes, where voltage-gated potassium channels cluster (Arancibia-Carcamo & Attwell, 2014; Boyle et al., 2001; Eshed et al., 2005; Rasband & Peles, 2016; Susuki et al., 2013). Together with the myelin sheath, the nodes of Ranvier maintain rapid conduction velocity along nerve fibers via saltatory conduction. Clustering of voltage-gated sodium channels in the node of Ranvier significantly enhances the efficiency of nerve conduction velocity (Harris & Attwell, 2012). Node and internode lengths can be fine-tuned to enhance conduction velocity in a rapid, energy-efficient manner (Arancibia-Carcamo et al., 2017; Ford et al., 2015).

Abnormal development or disruption of the nodal and paranodal microdomains contributes to a wide range of neurological diseases (Susuki, 2013). Disruption of the attachment between the terminal myelin loops and axolemma at the paranode results in mislocalization of voltage-gated ion channels across the microdomains. This mislocalization leads to lower action potential amplitude, slower conduction velocity, and reduced neuron excitability in mouse models of neurodegenerative diseases, such as multiple sclerosis and Charcot-Marie-Tooth disease (Crawford et al., 2009; Devaux & Scherer, 2005). While there is abundant information about molecular microdomains in the node of Ranvier in other nervous systems, the study of node microdomains in the peripheral auditory system is limited. Arrangement and maturation of voltage-gated ion channels are important for saltatory conduction in the axon initial segments and axonal nodes of the cochlea (Hossain et al., 2005; Kim & Rutherford, 2016). It is unknown how the nodes, with their special characteristics for fine-tuning conduction velocity, affect maturation of AN function and hearing onset. Using a multidisciplinary approach, we identified two distinct types of Ranvier nodes—the axonal node and the ganglion node—in the peripheral AN. We report the developmental and structural features of these two types of Ranvier nodes, and relate these to the spatiotemporal development of myelin. Our data identified the different structural characteristics of two cochlear glial cells, Schwann cells and satellite cells. Our data also aligned these cells with formation axonal nodes (Schwann cells) and ganglion nodes (satellite cells). Most importantly, comprehensive structure-function correlation analyses revealed that the length of the axonal node was associated with conduction speed and neural synchrony, whereas the length of the ganglion node was associated with amplitude growth rate. These data support that the different node types and their unique characteristics support different aspects of AN development and function.

Loss or dysfunction of the AN is a key cause of sensorineural hearing loss, the most common type of hearing loss. Extrinsic factors such as exposure to noise and aging can cause degeneration of AN fibers and spiral ganglion neurons (SGNs) (Spoendlin, 1984; Bao & Ohlemiller, 2010; Liberman, 2017; Liberman & Kujawa, 2017). Previous studies showed that an initial consequence of cochlear injury (such as noise exposure) is a loss of synaptic connections between sensory hair cells and the AN, which occurs before the degeneration in the other structural elements of the cochlea (Kujawa & Liberman, 2006; Lin et al., 2011; Liberman, 2015; Liberman & Kujawa, 2017). This finding highlighted the importance of AN degeneration in the etiology of sensorineural hearing loss. Approximately 90-95% of AN fibers are myelinated by cochlear glial cells. Emerging literature suggests that dysregulation of glial cells in the peripheral AN in several pathological conditions (including genetic defects, noise exposure and hair cell loss) is associated with hearing impairment (El-Badry et al., 2007; Jyothi et al., 2010; Akil et al., 2015; Kurioka et al., 2016; Panganiban et al., 2018; Rattay et al., 2013; Tang et al., 2006; Wan & Corfas, 2017). However, the link between the disruption of nodal structures and AN functional deficiency in sensorineural hearing loss is largely unknown.

Using an established mouse model of age-related hearing loss, we found structural abnormalities in the node of Ranvier in ANs of aged mice. In particular, we discovered changes in the satellite cell-associated ganglion node, implicating glial cell dysfunction as a mechanism of age-related hearing loss. The presence of these different node types, their unique structural and functional characteristics, and their changes that result from aging highlight the need to better understand nodal microstructure and its contribution to auditory function and disease.

## Results

### Myelination of the neuron soma is preceded by axonal myelination during postnatal AN development and hearing onset

The onset of hearing varies in mouse strains but generally occurs between postnatal day 9 (P9) and P14 (Alford & Ruben, 1963; Ehret, 1976; Song et al., 2006; Sonntag et al., 2009). In the CBA/CaJ mice used in this study, the external auditory meatus often opened around P11. To confirm the critical time points of postnatal development of AN and hearing onset in these mice, auditory brainstem response (ABR) was measured at P12, P14, P21 and 1 month (1M) (**Figure 1q, Supplementary Table 1**). Reliable ABRs with a clear wave I peak were recorded starting at P12. By P14, the ABR wave I thresholds improved approximately 25 to 30 dB across the frequencies tested. Threshold responses were significantly improved between each successive age group, except for P21 versus 1M for most frequencies tested, suggesting that hearing sensitivity reached maturity by P21. At P21 and 1M, ABR thresholds approximated those observed in young adult CBA/CaJ mice (Liu et al., 2019; Xing et al., 2012). These ABR data confirmed that the time points of P14, P21 and 1M constituted an appropriate developmental continuum to evaluate AN structure and its relationship to hearing onset.

**Figure 1.**
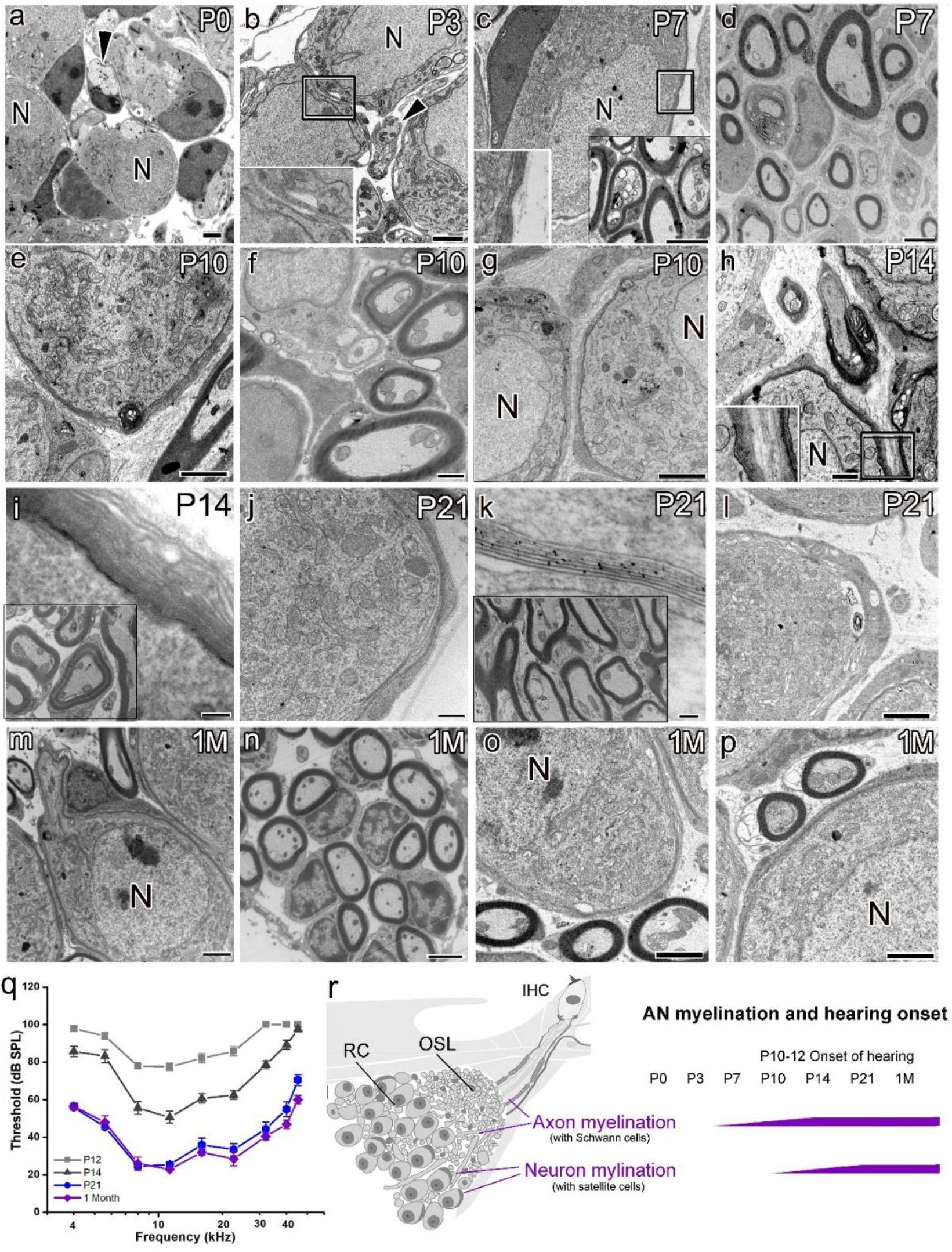
Myelination of spiral ganglion neurons occurs later than that of their axons during postnatal development of peripheral auditory nerve and hearing onset in mouse. (**a-p**) Transmission electron microscopy imaging showed distinct timing and structural features of myelination in neurons and their axons of the mouse peripheral auditory nerve (AN) at ages P0 (**a**), P3 (**b**), P7 (**c,d**), P10 (**e-g**), P14 (**h,i**), P21 (**k-m**) and 1 month (1M) (**m-p**). Unmyelinated axons (black arrowheads) and their neuron cell bodies were seen at P0 (**g**) and P3 (**h**). Enlargement of boxed area in **b** shows a thick, single layer of cytoplasm-filled satellite cell process encircling spiral ganglion neurons. Schwann cell-associated compact myelin sheets around AN axons were seen around P7 (The bottom-left boxed area in **c**). However, only 2-3 loose layers of satellite cell cytoplasmic processes enclosed the neurons (enlargement of the boxed area in **c**). Peak axonal myelination occurs between P7 and P10 (**c-f**), although the compact myelin sheets show variable in thickness of the enclosing axons of mixed caliber at P7 (**d**). By P14, axon myelination was nearly complete with compact myelin sheets (boxed area in **i**), similar to those axons seen at P21 (boxed area in **k**) and 1M (**m-p**). Increased layers of loose satellite cell process were seen around neuron somas from P10 (**e, g**) to P14 (enlargement of boxed area in **h**, **i**). The arrowhead in **g** highlighting the thick cytoplasm-filled process enwrapping the soma. Compact myelin sheets were seen around spiral ganglion neurons by P21 (**j,k,i**), similar to those at 1M (**m, o, p**). (**q**) Auditory brainstem response (ABR) wave I threshold significantly improved from P12 to P21 (mixed-effects model, p<0.0001). Responses between P21 and 1M were largely unchanged for most frequencies, except for the significantly different responses at 22.6, 40, and 45.2 kHz (mixed-effects model with Benjamini-Hochberg post-hoc analysis, see **Supplementary Table 1**). Error bars indicate standard error of the mean in **q**. (**r**) Schematics showing myelinated nerve fibers passing through the osseous spiral lamina (OSL) to innervate inner hair cells (IHC), and the timing of myelination across postnatal ages highlighting the delay of myelination of spiral ganglion somata compared to earlier myelination of axons. RC, Rosenthal’s canal.

To better understand how nodal structures form in the AN, we examined ultrastructural features of myelination in the cochleas of CBA/CaJ mice at stages before hearing onset (P0, P3, P7, and P10) and after hearing onset (P14, P21, and 1M; **Figure 1q**) using transmission electron microscopy (TEM) (**Figure 1**). Myelin sheets formed around the AN axons at P6 and older stages (**Figure 1c-f,h,k,m-p**), but not at P0 (**Figure 1a**) or P3 (**Figure 1b**). By P7, Schwann-cell processes surrounded axons with myelin sheets in a variety of thicknesses (**Figure 1d**). Variation in myelin thickness diminished after P10. By P21 (**Figure 1k**), myelination was mostly complete, with myelin sheath thickness similar to that observed in 1M-old mice (**Figure 1n**) and in young adult mice reported previously (Lang et al., 2011; Xing et al., 2012). Myelination of SGN somas began at P10, with 2 to 3 layers of thick, cytoplasm-filled satellite-cell processes seen around the neuron cell bodies (**Figure 1e,g**). By P14, the layers of satellite cell processes were increased but no compact myelin sheets formed (**Figure 1i**). Compact myelin sheets were first detected around neurons at P21 and 1M (**Figure 1j-m,1o,1p**). These data revealed that 1) myelination of AN axons by Schwann cells begins around P7 and finishes around P14; 2) myelination of SGN soma by satellite cells, another type of glial cell in the cochlea, starts at P10, shortly before hearing onset and completes by P21; 3) an approximately 7-day delay occurs between formation of the compact myelin sheet around axons (by Schwann cells) and around the somata (by satellite cells); and 4) at least two types of cochlear glial cells are associated with AN myelination in mouse cochlea (**Figure 1r**).

### Expression profiles of genes composing nodal structural microdomains exhibit special temporal patterns in the developing AN

To characterize nodal assembly processes during postnatal AN development and hearing onset, we examined the expression profiles of myelin- and node-related genes at several critical developmental time points of myelination, including P7, P14, and P21. These timepoints were identified in our TEM studies (**Figure 1**). P3 was also selected as a control for the pre-myelinated AN. RNA-sequencing (RNA-seq analyses) showed that 1502 of 1727 myelin-related genes were significantly regulated (p-adjusted <0.05 for at least one pairwise comparison) (**Figure 2a, Supplementary Table 2**). Additionally, 38 of the 42 genes identified as node-related were differentially expressed during this developmental period (**Supplementary Table 3**). Hierarchical clustering of expression data for the node-related genes revealed that the genes were separable into three different clusters of temporal expression profile (**Figure 2b**). Cluster 1 (peak expression P3-P7) contained genes important for onset of nodal microdomain formation, including these affecting cell-cell interaction, scaffolding, cell adhesion, and extracellular matrix (Custer et al., 2003; Ervasti & Campbell, 1993; Woods et al., 1996). Cluster 1 also contained *Scn2a,* which encodes the voltage-gated sodium channel Na_v_1.2. This channels is a marker of immature nodes of Ranvier later replaced by Na_v_1.6 (encoded by *Scn8a)* in adult nodes of Ranvier after the onset of myelination (Kaplan et al., 2001). Cluster 2 (peak expression at P7-P21) contains genes involved in the onset of paranodal junction assembly (Eylar et al., 1971; Ishibashi et al., 2004; Kwon et al., 2013; Li et al., 1994; Orthmann-Murphy et al., 2009; Quarles et al., 1973; Snipes et al., 1992; Zonta et al., 2008). This cluster includes *Cntn1*, which encodes the Cntn1 axo-glial connector protein that attaches axolemma and myelin lamellae at the paranode (Boyle et al., 2001). Cluster 3 (peak expression at P14-21) is comprised of genes encoding structural proteins of the electrically excitable nodal domain (Huang et al., 2017; Lacas-Gervais, 2004), and axo-glial connector proteins at the paranode (Rios et al., 2003) and juxtaparanode (Poliak et al., 1999; Poliak & Peles, 2003). Cluster 3 also included genes encoding voltage-gated potassium channels of the juxtaparanode (Rasband et al., 1998; M N Rasband & Trimmer, 2001) and the mature nodal Na_v_1.6. Based on these findings, node formation begins around P3, paranodal domain formation and myelination around P7, and juxtaparanode assembly around P14. The time-dependent up-regulation of these genes encoding nodal structural proteins and voltage-gated ion channels shows that nodal structure develops before hearing onset and has fully formed microdomains by the time of hearing onset.

**Figure 2.**
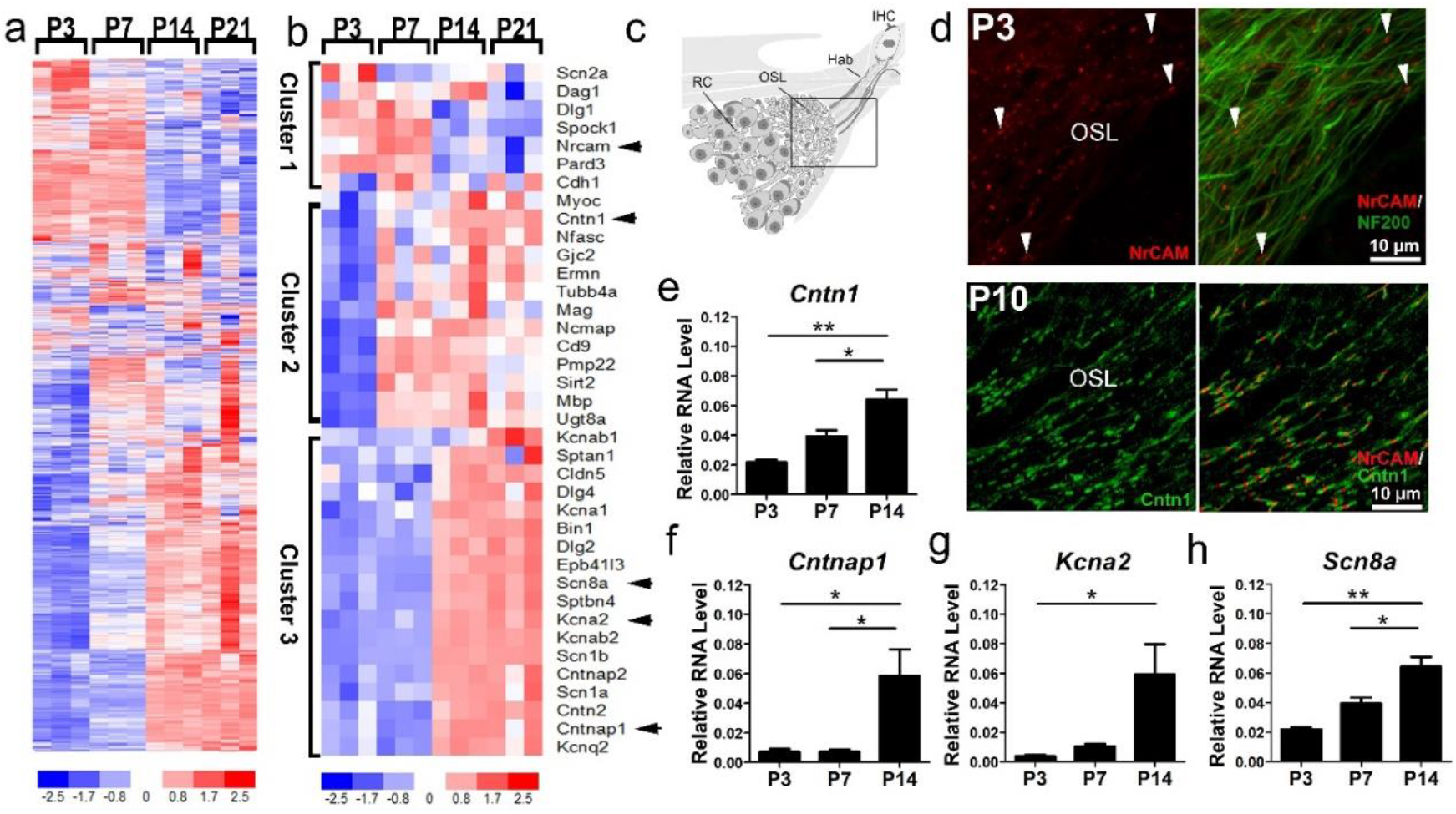
Expression profiles of genes composing nodal structural domains exhibit temporal clustering. (**a**) Heatmap of myelin-related genes differentially expressed in the AN during postnatal development. Of the 1727 myelin-related genes, 1502 were significantly different (p-adjusted <0.05) for at least one of the pairwise comparisons (**Supplementary Table 2**). Myelin-related genes used in this analysis were compiled in our previous study (Panganiban et al., 2018). (**b**) Heatmap of node-related genes differentially expressed in AN during postnatal development. Of 42 node-related genes, 38 were significantly different (p-adjusted <0.05) for at least one pairwise comparison (**Supplementary Table 3**). Genes were segregated into three different clusters based on temporal expression profile. (**c**) Schematic of the AN. Box indicates the area of immunostaining in **d** (top panels). Immunostaining shows the presence of nodal NrCAM along axonal processes labeled by NF200 as early as P3 (**d**, bottom panels). Immunostaining shows an abundance of axonal nodes with paranodal Cntn1 flanking nodal NrCAM (**e-h**). qPCR experiments validate the presence of nodal (**h**), paranodal (**e,f**), and juxtaparanodal (**g**) genes in a separate set of P3, P7, and P14 ANs. ANOVAs with Bonferroni corrections for multiple comparisons were performed to show differential expression among the time points (* = p<0.05, ** = p<0.01); n = 3 mice per age group.

As shown in **Figure 2d-h**, our RNA-seq findings were validated by immunostaining and reverse transcription quantitative polymerase chain reaction (RT-qPCR). NrCAM node protein, which is encoded by *Nrcam* and responsible for initiating node formation (Custer et al., 2003), was detected in ANs at P3 and P10 (**Figure 1d**). Dual-immunostaining analysis with NrCAM and neuronal marker NF200 showed that NrCAM is present very early in nodes of the mouse AN (**Top panels of Figure d**). At P3, NrCAM was present only in the hook area of the AN, which is the end of the basal point of the cochlea turn. This may indicate that nodal assembly occurs in a basal to apical gradient, which would be consistent with the basal to apical development pattern of other cochlear structures. Dual-immunostaining analysis with NrCAM and the paranodal protein Cntn1 revealed that distinct nodal (NrCAM+) and paranodal (Cntn1+) microdomains were present at P10 (**Bottom panels of Figure 1d**). With RT-qPCR analysis, we assessed the temporal expression pattern of node-related genes in Cluster 2 (*Cntn1*; **Figure 2e**) and Cluster 3 (*Cntnap1, Kcna2,* and *Scn8a*; **Figure 2f-h**). Our statistical analysis confirmed that all genes were significantly upregulated between P3 and P14 (one-way ANOVA with Bonferroni post hoc adjustment; **Table 1**).

**Table 1.** One-way ANOVA with Bonferroni’s multiple comparisons correction of relative RNA concentrations of key nodal genes across postnatal ages

| Gene | ANOVA<br>p value | DF<sub>n</sub>,<br>DF<sub>d</sub> | F | Comparisons | Mean diff. | 95% CI | Adjusted *p*<br>value |
| --- | --- | --- | --- | --- | --- | --- | --- |
| *Cntn1* | 0.002 | 2,6 | 23.060 | P3 vs. P7 | -0.018 | -0.038 to 0.003 | ns |
|  |  |  |  | P3 vs. P14 | -0.043 | -0.063 to -0.022 | ≤ 0.05 |
|  |  |  |  | P7 vs. P14 | -0.025 | -0.046 to -0.004 | ≤ 0.01 |
| *Cntnap1* | 0.02 | 2,6 | 8.048 | P3 vs. P7 | -0.0001 | -0.049 to 0.049 | ns |
|  |  |  |  | P3 vs. P14 | -0.051 | -0.100 to -0.003 | ≤ 0.05 |
|  |  |  |  | P7 vs. P14 | -0.051 | -0.100 to -0.003 | ≤ 0.05 |
| *Kcna2* | 0.03 | 2,6 | 6.634 | P3 vs. P7 | -0.007 | -0.062 to 0.048 | ns |
|  |  |  |  | P3 vs. P14 | -0.056 | -0.110 to -0.001 | ≤ 0.05 |
|  |  |  |  | P7 vs. P14 | -0.049 | -0.100 to 0.006 | ns |
| *Scn8a* | 0.002 | 2,6 | 23.060 | P3 vs. P7 | -0.018 | -0.038 to 0.003 | ns |
|  |  |  |  | P3 vs. P14 | -0.043 | -0.063 to -0.022 | ≤ 0.01 |
|  |  |  |  | P7 vs. P14 | -0.025 | -0.046 to -0.004 | ≤ 0.05 |

### Heminodal clustering in the postnatal AN occurs in a spatially and temporally dependent manner

In the central nervous system, action potentials are generated at the axon initial segment, a site located between the neuron cell body and the axon (Palay et al., 1968). However, the generation of action potential in the peripheral AN is not fully understood. The inner hair cells (IHCs) are innervated by the peripheral processes of SGNs, whose bipolar cell bodies are located within Rosenthal’s canal (**Figure 1r**; **Left panel of Figure 4**). Several spike-generating Na_v_ and K_v_ channel proteins have been identified at multiple sites along the peripheral AN (Hossain et al., 2005; Kim & Rutherford, 2016). These sites include nodal structures in the peripheral axons and the heminodes.

Heminodes are a specialized form of the axon initial segment and are located on type I SGN peripheral processes where the distal osseous spiral lamina (OSL) and the habenula are 20-40 μm from the presynaptic ribbon-type active zone on an IHC (Hossain et al., 2005; Liberman, 1980). A previous study using computational modeling of patch-clamp recording from rodent ANs revealed that action potentials of the AN begin in these heminodes (Rutherford et al., 2012). To study heminodes formation in postnatal cochleas, immunostaining was conducted on whole-mount AN tissue preparations from mice at ages P7, P10, P14, and 1M for the nodal marker NrCAM or voltage-gated Na_v_1.6 channel protein, together with the paranodal marker Cntn1 (**Figure 3**). Our data showed that before P14, the molecular components of the heminodes (as marked by NrCAM or Nav1.6) are still positioned towards the habenula (**Figure 3a-a’’,b-b’’,e,f**). This is less apparent in the basal turn at P7 (**Figure 3a”**), as more NrCAM+ heminodes are clustered underneath the IHCs. While several heminodes visibly approach from the OSL region in the basal turn at P7, heminodes are almost completely clustered under IHCs in the basal turn at P10 (**Figure 3b**”). The number of unclustered heminodes approaching the habenula from the OSL is even more apparent in the middle turn at P7 to P10 (**Figure 3a’,b’**) and in the apical turn at P7 to P14 (**Figure 3a,b,c**). By P14, the close clustering of heminodes underneath IHCs is mostly complete in the basal and middle turns. In ANs; of 1M young-adult mice, heminodal clustering is complete in all three turns (**Figure 3d,d’,d**”). These observations support that heminode migration and clustering underneath the IHCs progresses in a basal to apical direction. The flanking Cntn1+ paranode also shows the position of the nodal component toward the habenula during P7 to P14 and highlights the distance between the heminode clusters and first axonal node. As shown in **Figure 3e-h**, the heminode clustering completes in the middle turn by P14, as 1) the clusters are directly under the IHCs and 2) the distance between the heminodes and the first axonal nodes are much longer (compared to that of P7) across the OSL.

**Figure 3.**
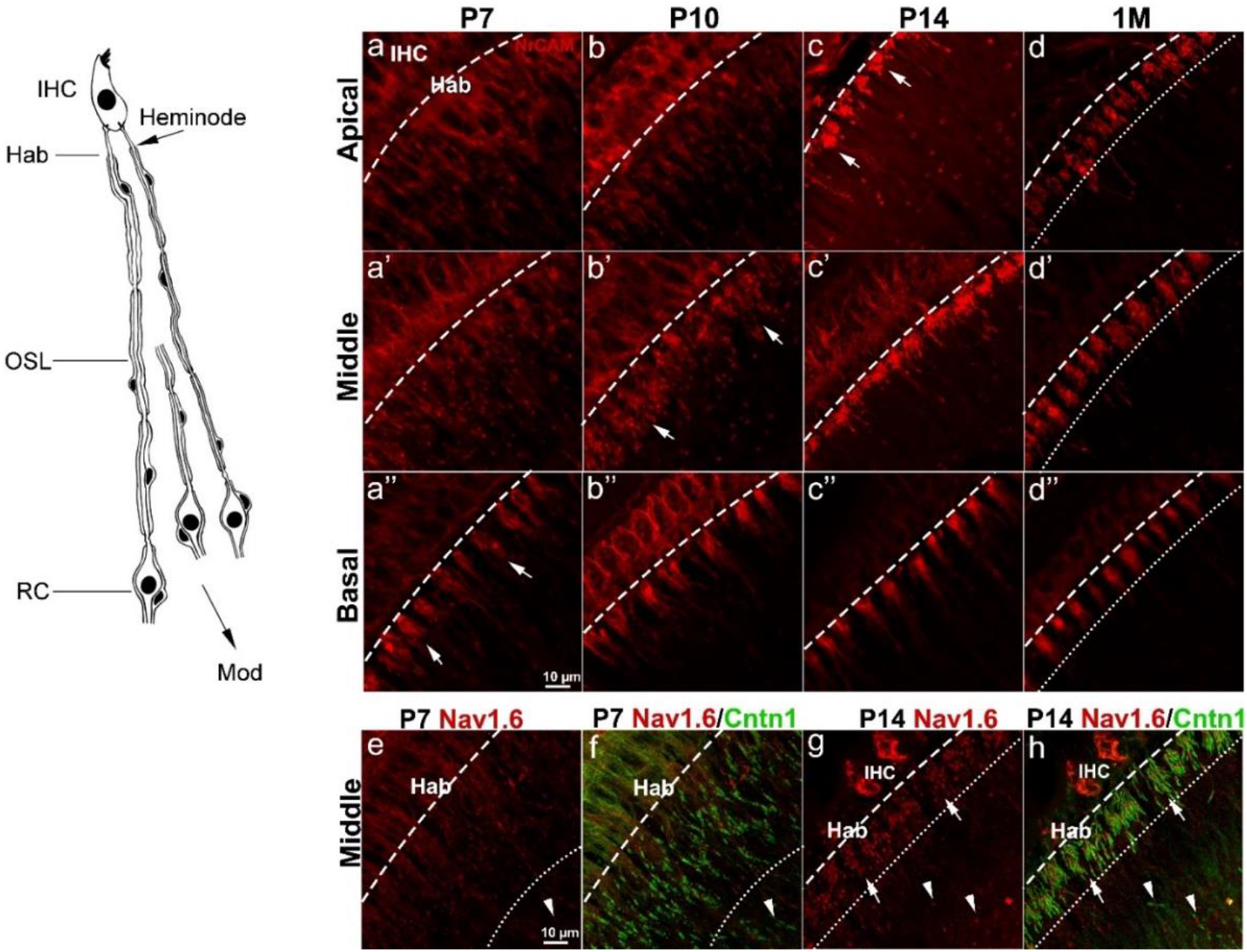
Clustering of AN heminodes across critical in postnatal development. The left panel is a schematic of peripheral AN fibers under an IHC illustrating the location of heminodes in the habenula (Hab) and the distal site of the osseous spiral lamina (OSL). (**a-c**) Progression of heminodal clustering indicated by NrCAM immunostaining is shown horizontally across critical time points of myelination (at P7, P10, P14, and 1M) and vertically across the location from the apical (**a-d**), middle (**a’-d’**), and basal turn (**a”-d”**). Arrows in **a”, b’**, and **c** indicate the organized clustering of NrCAM+ heminodes of each fiber unit condensed underneath the IHCs which progress in a basal-to-apical manner. Immunostaining of 1M ANs in **d-d”** highlights the canonical clustering of heminodes of all three cochlear turns (**e-f**). Dual-immunostaining of nodal Nav1.6 (red) with paranodal Cntn1 marker (green) shows progression of heminodal clustering of spike-generating ion channels in the AN from P7 (**e,f**) to P14 (**g,h**). Arrows in **g,h** highlight the completed clustering of heminodes. Arrowheads in **e-h** indicate the location of the first axonal nodes after the heminodal clusters. Dashed lines indicate the location of habenular openings along the whole-mounts. Dotted lines indicate the space between the habenular opening and the heminodes most distal to IHCs. RC, Rosenthal’s canal; Mod, modiolus. Scale bars: 10 μm in **a’’** (for **a-d’’**); 10 μm in **e** (for **e-h**).

### Assembly patterns of nodal and paranodal proteins reveal the formation of two distinct types of the nodal structures in the AN

To study nodal structure formation in postnatal ANs, we performed dual-immunostaining for the nodal marker NrCAM and the paranodal marker Cntn1 on frozen sections prepared from cochleas at P7, P10, and P14 (**Figure 4**). Results showed that heminodes are only found in the habenular region (**Figure 4a,e,i**), corroborating the our findings in whole-mount preparations of AN (**Figure 3)**. Along SGNs running through the OSL, Rosenthal’s canal and central modiolus region (Mod), we identified two types of nodes of Ranvier: 1) axonal nodes, found along the axon in the peripheral side of AN fibers at the OSL and the central side of AN fibers at the Mod (**Figure 4a,e,I,b,f,j,c,g,k)**; and 2) ganglion nodes, found in Rosenthal’s canal and flanking type I SGN somata (**Figure b,f,j,d,h,l)**. At P7, axonal nodes (NrCAM+), are present and appear to be flanked by two partially formed paranodes (Cntn1+) (**Figure 4c**). In contrast, most ganglion nodes do not have Cntn1 immunoreactivity (**Figure 4d**). By P10, axonal nodes express Cntn1 on both paranodal domain positions (**Figure 4g**). Cntn1 activity was also seen in the axonal paranodal position but not in the paranode position on the somal side of the ganglion nodes (**Figure 4h**). By P14, we saw distinct immunostaining patterns of nodal (Cntn 1+) and paranodal (NrCAM+) elements in the axonal node and ganglion node (**Figure 4k,l**). This staining pattern was maintained in cochleas of young-adult mice (**Figure 5m,n)** and aged mice **(Figure 8a)**, and was also seen in aged human temporal bone **(Figure 8c)**. Taken together, these data demonstrated distinct features in two types of nodal structures with unique features in the AN: 1) there are two Cntn1+ paranodal elements in the axonal node, whereas the ganglion node contains only one Cntn1+ paranodal element; and 2) the NrCAM+ nodes is significantly shorter in axonal nodes than ganglion nodes (unpaired t-tests, p< 0.0001) (**Figure 6** and **Table 2**). The difference in nodal structural was also occurred in all reported axonal node lengths when compared to corresponding ganglion node lengths from the same age group and cochlear turn (e.g., P7 apical axonal node vs P7 apical ganglion node). In addition, delayed development of the ganglion node corresponds with a similar delay in myelination of soma, as compared to peripheral axons (**Figure 1**).

**Table 2.** Measurements of axonal and ganglion node length

| Mean axonal node length (μm) |  |  |  |  |  |  |
| --- | --- | --- | --- | --- | --- | --- |
|  | Apical |  | Middle |  | Basal |  |
| P7 | 2.08 ± 0.11 |  | 2.01 ± 0.10 |  | 1.97 ± 0.07 |  |
| P10 | 1.69 ± 0.13 |  | 1.60 ± 0.12 |  | 1.67 ± 0.20 |  |
| P14 | 1.60 ± 0.10 |  | 1.46 ± 0.09 |  | 1.47 ± 0.09 |  |
| P21 | 1.69 ± 0.14 |  | 1.66 ± 0.08 |  | 1.68 ± 0.09 |  |
| 1M | 1.91 ± 0.25 |  | 1.95 ± 0.19 |  | 1.98 ± 0.11 |  |
| Mean ganglion node length (μm) |  |  |  |  |  |  |
|  | Apical |  | Middle |  | Basal |  |
| P7 | 3.86 ± 0.65 |  | 3.94 ± 0.22 |  | 3.84 ± 0.64 |  |
| P10 | 3.98 ± 0.58 |  | 3.65 ± 0.34 |  | 3.79 ± 0.29 |  |
| P14 | 3.81 ± 0.28 |  | 3.38 ± 0.20 |  | 3.40 ± 0.17 |  |
| P21 | 3.36 ± 0.16 |  | 3.14 ± 0.10 |  | 2.79 ± 0.23 |  |
| 1M | 3.34 ± 0.22 |  | 2.88 ± 0.27 |  | 3.20 ± 0.18 |  |
| Mean difference between axonal and ganglion node length (μm) by cochlear turn and age |  |  |  |  |  |  |
|  | Difference<br>Apical | SE | Difference<br>Middle | SE | Difference<br>Basal | SE |
| P7 | -1.78<sup>†</sup> | 0.27 | -1.93<sup>‡</sup> | 0.10 | -1.87<sup>†</sup> | 0.26 |
| P10 | -2.28<sup>†</sup> | 0.24 | -2.06<sup>‡</sup> | 0.15 | -2.12<sup>‡</sup> | 0.14 |
| P14 | -2.21<sup>‡</sup> | 0.11 | -1.91<sup>‡</sup> | 0.08 | -1.93<sup>‡</sup> | 0.07 |
| P21 | -1.67<sup>‡</sup> | 0.08 | -1.48<sup>‡</sup> | 0.05 | -1.10<sup>‡</sup> | 0.09 |
| 1M | -1.43<sup>‡</sup> | 0.13 | -0.93<sup>†</sup> | 0.12 | -1.22<sup>‡</sup> | 0.08 |

Unpaired t-test, <sup>†</sup> $p < 0.0001$ , <sup>‡</sup> $p < 0.00001$ . SE, standard error.

**Figure 4.**
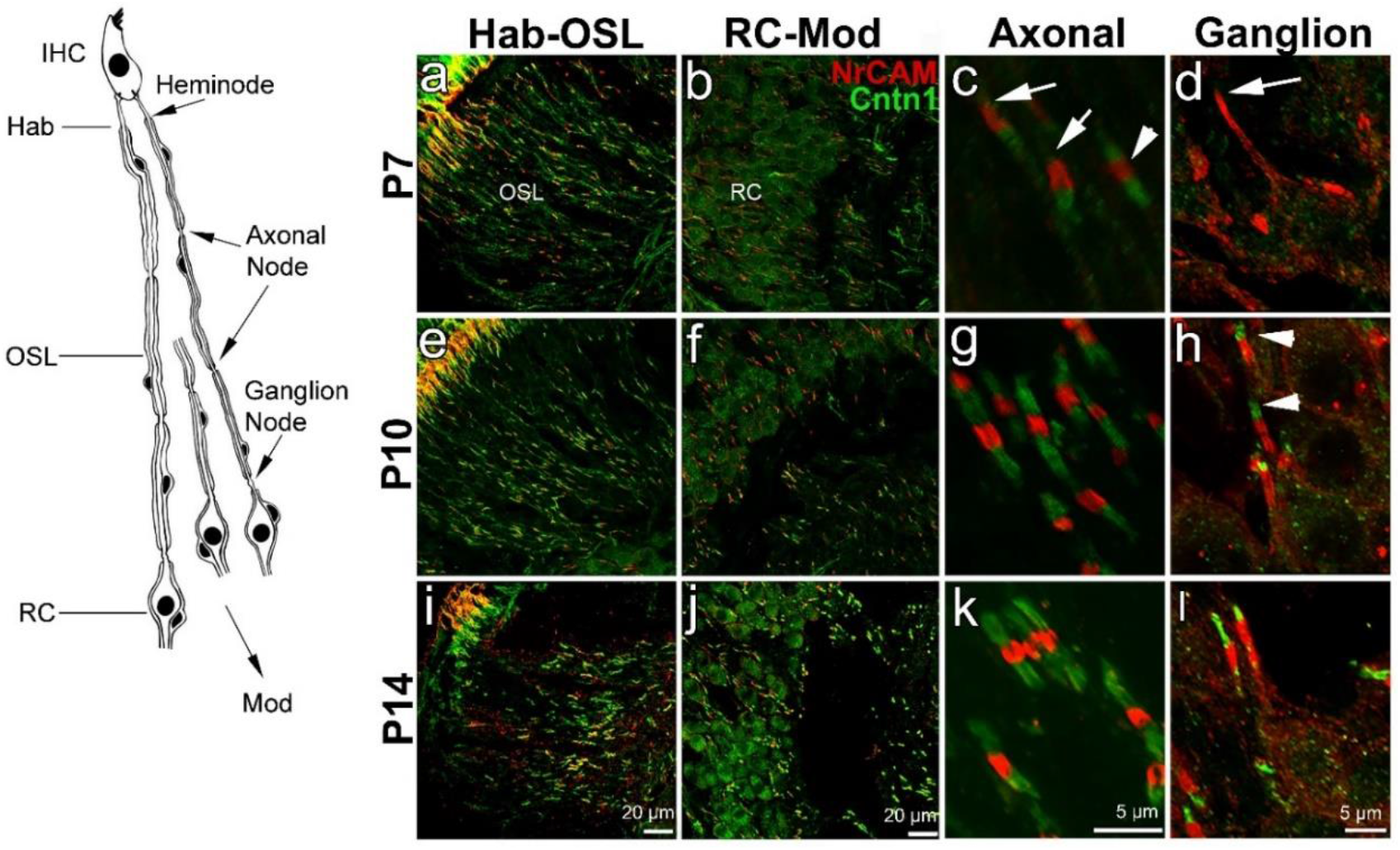
Assembly of nodal and paranodal proteins comprising the axonal and ganglion nodes in the postnatal AN. Left panel is a schematic of peripheral AN fibers illustrating the locations of heminodes in the habenula (Hab), the distal site of the osseous lamina (OSL), axon nodes within the OSL and ganglion nodes within Rosenthal’s Canal (RC) (**a-h**). Panels show the assembly of the nodes, marked by NrCAM (red), and paranodes, marked by Cntn1 (green) from P7, P10, and P14. **a,e,i** show heminodes and axonal nodes at the Hab and OSL, respectively. **b,f,j** show ganglion nodes located in the RC and the axonal nodes passing through the RC to the modiolus. **c,g,k** are enlargements of axonal nodes found in the OSL in P7 (**c**), P10 (**g**), and P14 (**k**). White arrows in **c** indicate axonal nodes with missing undeveloped paranode flanks and the white arrowhead indicates a node completely flanked by two paranodes on either side. **d,h,l** are enlargements of ganglion nodes found in the RC at P7 (**d**), P10 (**h**), and P14 (**l**). White arrow in **d** indicates a ganglion node without a Cntn1-marked paranodal flank. White arrowheads in **h** indicate paranode flanks of the ganglion nodes. Images were taken at the middle turn. Scale bars: 20 μm in **i** (for **a,e,i**); 10 μm in **j** (for **b,f,j**); 5 μm in **k** (for **c,g,k**); 5 μm in **l** (for **d,h,l**).

**Figure 5.**
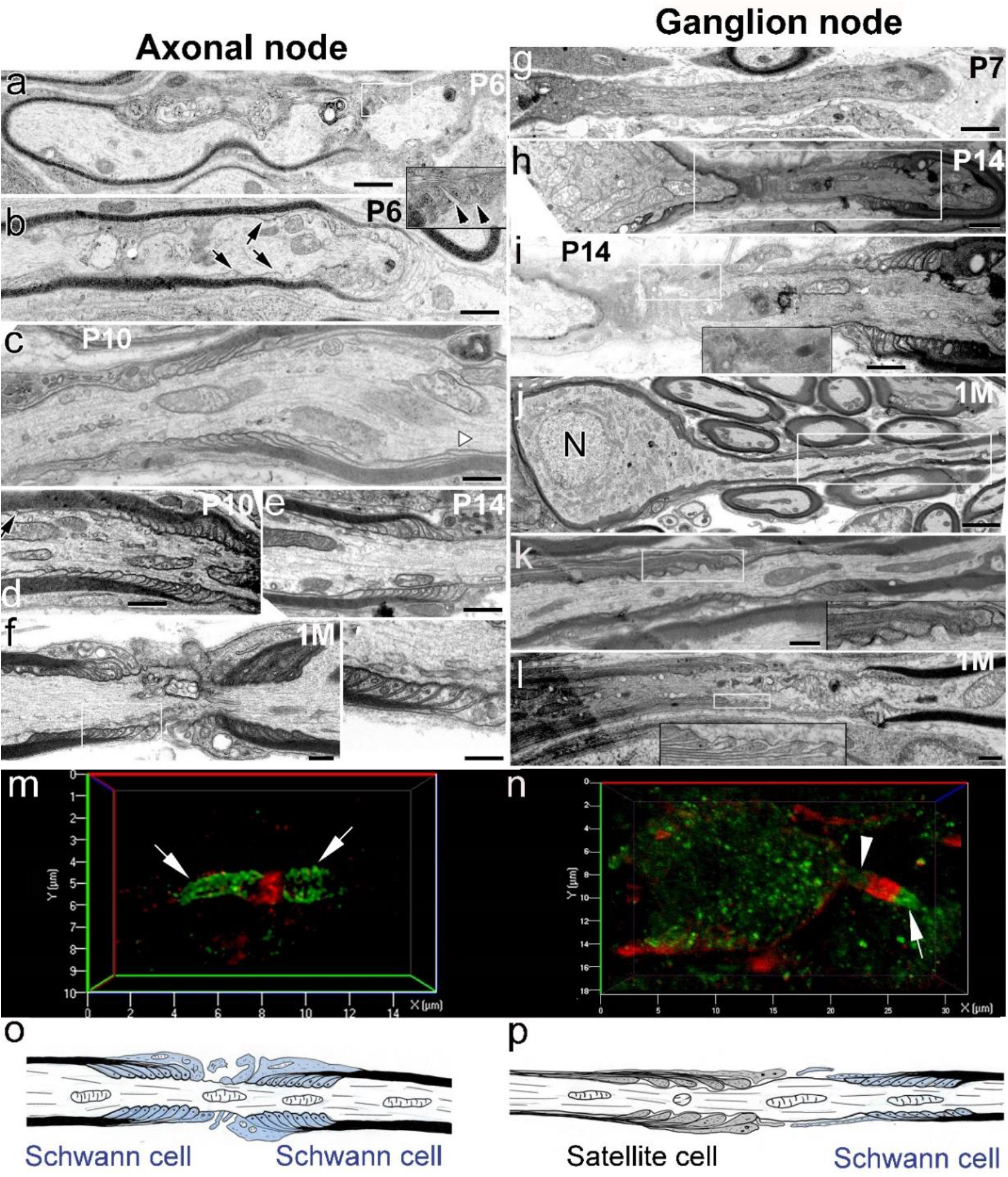
Transmission electron microscopy revealing distinct paranodal ultrastructure in the axonal and ganglion nodes of the peripheral AN. (**a-f**) Myelin terminal loops at both sides of the axon node assembly by Schwann cells at the paranodal domain started forming around P6 but were not completed until P10. Terminal loop layers were non-compact (black arrowheads in the enlargement of the white-line-framed boxed area in **a)** and several terminal loops are still migrating from the intermodal region (black arrows in **b)**. Paranodal domain assembly is mostly complete by P10 (**c-d**). A white arrowhead in **c** and a black arrow in the upper-left corner in **d** show the terminal loops in the juxtaparanodal domain. Complete, compact organization of terminal loops at both paranodal domains of the axonal node at P14 (**e**) is similar to the arrangement seen in young adult mice (1M) (**f**, left panel is an enlargement of boxed area). (**g-l**) Delayed formation and distinct structural of the paranode (formed by satellite cells) in the somal side of the ganglion node compared to those in the axonal side of the node. At P7, the ganglion paranode was not identified (**g**). By P14, a myelin terminal loop-like structure was seen in the somal side of the ganglion node, whereas myelin terminal loops were completely formed in the axonal side of the ganglion node (**h-i**). The image in **i** is an enlargement of boxed areas in **h**. Image at the bottom of **i** is an enlargement of boxed area in **I**. Complete assembly of the ganglion node at both sides of the node by satellite cells and Schwann cells occurs by 1M (**j-l**). Enlargements of boxed areas in **k,l** show loose satellite cell terminal loops at the ganglion paranodal domain. The white boxes indicate the location of enlargements in the black boxes (**m,n**). Dual immunostaining with paranodal marker (NrCAM, red) and nodal maker (Cntn1, green) showed a different canonical structure of axonal (**m**) and ganglion (**n**) nodes in the AN of a young adult mouse. A super-resolution image of an axonal node reveals its terminal myelin loops (Cntn1+; green). NrCAM+ paranodal element (arrow) is present at both sides of the axonal node (**m**) but only one side (axon side) of the ganglion node (**n**). An arrowhead indicates no NrCAM+ paranodal element at the somal side of the ganglion node (**o,p**). Schematics illustrate the distinct structural patterns and glial components of the axonal (**o**) and ganglion (**p**) nodes. Paranode domains of the axonal node contain two similar clusters of tightly-assembled terminal loops provided by two adjacent Schwann cells. A cluster of loose myelin terminal loops from a satellite cell (somal side) and a cluster of tightly-assembled terminal loops provided by a Schwann cell (axonal side) make up the two paranodal domains of the ganglion node. Scale bars: 400 nm in **a-f,h,I,k,l**; 150 nm in left panel in **f**; 800 nm in **g**; 2 μm in **j**.

**Figure 6.**
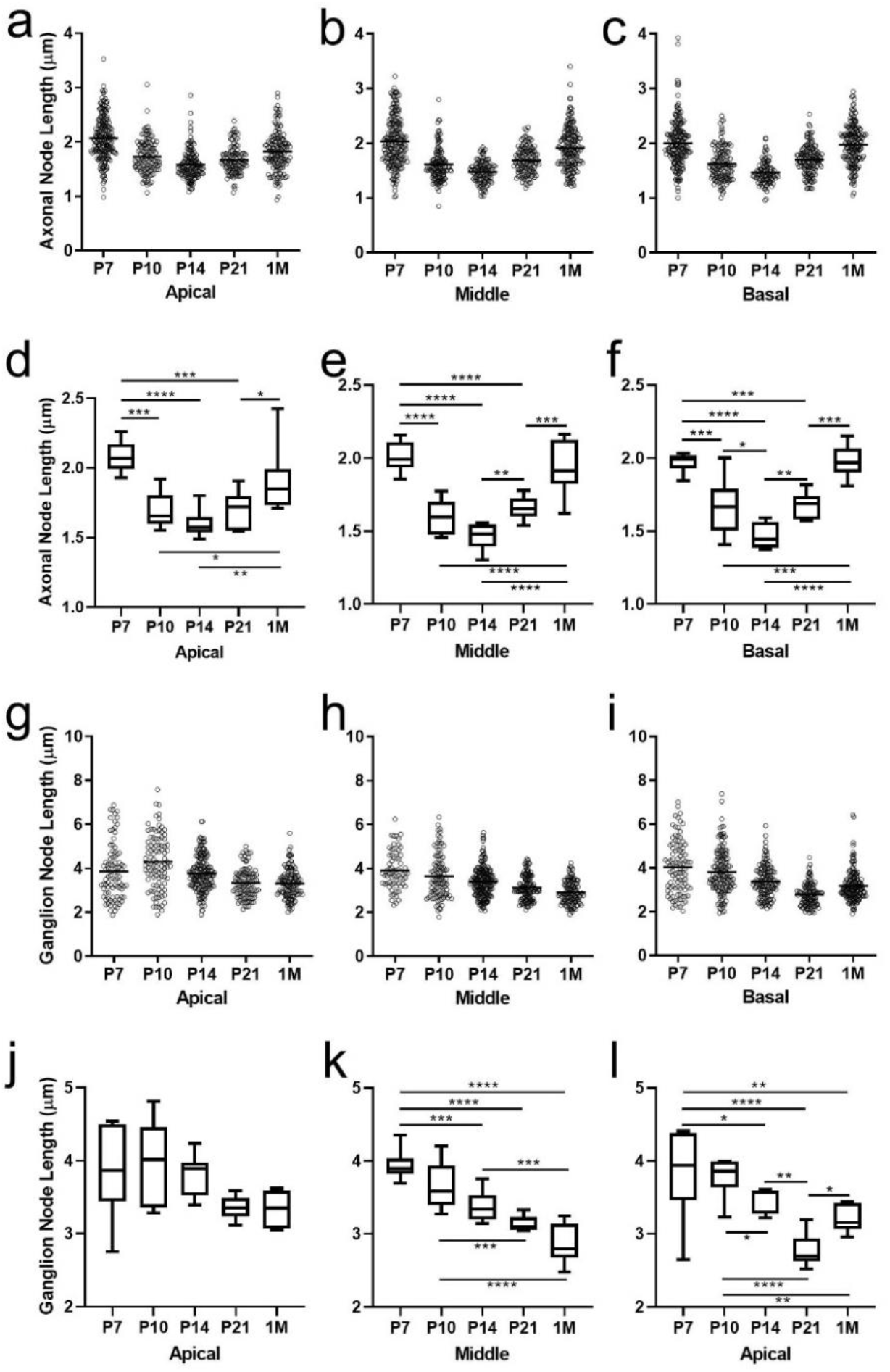
Axonal and ganglion node lengths change between critical developmental time points. Axonal and ganglion node length measurements from ANs were taken from the apical, middle, and basal cochlear turns at P7, P10, P14, P21, and 1M. Scatter plots show all length measurements taken of axonal (**a–c**) and ganglion (**g–i**) nodes in mice from each age group. Box plots show the average length of axonal (**d–f**) and ganglion (**j–l**) nodes in mice from each age group. One-way ANOVA and post-hoc tests, done using Benjamini-Hochberg method to control for false discovery rate (Q = 0.05), were performed on **d-f** and **j-l**. Bars and asterisks indicate pairwise comparisons with significant discoveries; * = q<0.05, ** = q<0.01, *** = q<0.001, **** = q<0.0001. nP7 = 6, nP10 = 6, nP14 = 8, nP21 = 7, n1M = 8. Average nodal lengths and statistical test results are shown in **Table 2**.

### Axonal and ganglion nodes in the postnatal AN have different connection patterns of axon-Schwann cells and axon-satellite cells in their paranodal microdomains

Axonal paranodes are generated by the terminus of the myelin lamellae from myelinating Schwann cells at the paranodal axolemma. Unlike the thin, nearly cytoplasm-free myelin lamellae ensheathing the internodes, the terminating lamellar loops at the paranode are filled with cytoplasm. These loops provide a larger surface area for axo-glial connections and glial-glial connections between the adjoining lamellae and terminal loops (see reviews by Poliak & Peles, 2003; Sherman & Brophy, 2005; **Figure 5f,l,o,p**). This axo-glial connection is accomplished by cell adhesion molecules, such as Cntn1. The terminal myelin loops and their axo-glial connections serve as physical barriers that separate the voltage-gated sodium channels in the node from the voltage-gated potassium channels in the juxtaparanode. These structures are critical for generating and propagating action potentials (Boyle et al., 2001; Rasband et al., 1998; Rios et al., 2000). To better understand the differences in 1) the structure of the two nodal types and 2) the axo-glial interactions in the axonal paranodes (formed by Schwann cells) and ganglion paranodes (formed by satellite cells), we analyzed the ultrastructural features of peripheral ANs in postnatal mice (**Figure 5**). At P6 and P10, terminal myelin loops in the axonal paranodes are still migrating to the paranodal domain from the internodal region (**Figure 5a-c**). By P14, the canonical paranodes are formed, showing terminal myelin loop heads connected to the paranodal axolemma and juxtaposed closely with no gaps, similar to those seen in young adult cochleas (**Figure 5f**).

As shown in **Figure 5g-I**, the terminal loops of the myelin lamellae from satellite cell were not found at the somal side of ganglion nodes until P14 (**Figure 5g-j**). At P14, the paranodal flank generated by a Schwann cell shows clear organization and a tight connection between the terminal myelin loops and the axolemma (**Figure 5i**). In contrast, the paranodal region generated by a satellite cell in the ganglion node showed no or incomplete attachment of the lamellae with the paranodal axolemma (**Figure 5i**). Satellite cell-lamellae are very few and are loosely attached to the axolemma. By 1M, the number of satellite cell-lamellae increased and the length of unmyelinated node gap between myelinated paranodes decreased compared to P14 (**Figure 5j-l**). However, the terminal loops of satellite cell appear only loosely connected to the paranodal axolemma (**Figure 5f,l**) when compared to the tightly connected terminal loops of Schwann cell (**Figure 5f**). Regardless of age of the auditor nerve, the ultrastructural differences were clearly evident between axon-Schwann cell and axon-satellite cell connections. Fewer myelin lamellae from satellite cells surrounded the soma and terminated at the ganglion paranode compared to Schwann cells. Also, high-resolution imaging analysis of the axonal node and ganglion node showed that Cntn1 was not expressed at the axon side of the ganglion node, which may contribute to the weaker connection between axons and satellite cells **(Figure 5m-p)**. The terminal myelin loop heads, though connected to the axolemma, have much more space between the loops and axolemma, suggesting that axon-satellite connections are weaker than axon-Schwann cell connections (**Figure 5o,p)**.

### Ganglion and axon node lengths in the AN are distinctive during postnatal development

As shown in **Figure 5f,j-l**, the ganglion node is longer than the axon node in ANs of young adult mice. To quantify and characterize how these nodes change during development, we measured the axonal (**Figure 6a-f**) and ganglion (**Figure 6g-l**) node lengths across the cochlear turns at pre-hearing onset (P7, P10) and post-hearing onset (P14, P21 and 1M) (**Table 2**). With one-way ANOVA with correction for multiple testing (Benjamini-Hochberg), we found that axonal nodes are significantly longer at P7 compared to P10 and P14 for all cochlear turns (**Figure 6a-f**). For example, in the middle turn, P7 axonal nodes are approximately 0.4 μm longer than P10 axonal nodes (p<0.0001) and 0.5 μm longer than P14 axonal nodes (p<0.0001).This may be due to the relatively large distance between two opposing myelin sheaths. As myelination continues, the opposing myelin sheaths formed by two Schwann cells move closer, forming the unmyelinated nodal gap. At P14, the axonal node gap is at its shortest for all three turns. By 1M, however, the axonal nodes have significantly increased in length compared to P14. The mean node length at P14 was ~1.5 μm, while at 1M the mean node length had increased to ~1.9 μm across all three turns.

An evaluation of how axonal node length changed during development revealed a dynamic pattern that was evident in all three cochlear turns. Nodes were longer at P7, shortened at P14, and then lengthened again at 1M (**Figure 6a-f**). Plotted graphically, the temporal profile creates a “V” shape. For the ganglion nodes, the mean length was significantly shorter at P21 than at P7 and P10 (mean diffs. P7 vs P21 = 0.80 μm, p<0.0001; P10 vs P21 = 0.51 μm, p = 0.0009), but was not significantly different from P14 (mean diff. P14 vs P21 = 0.24, p = 0.06 for the middle turn (**Figure 6h,k**). In the middle turn, the ganglion node lengths continually shortened with age, from P7 towards 1M (**Figure 6h,k**). In the basal turn, the ganglion node continually shortened from P7 to P14, but unexpectedly lengthened again at 1M (**Figure 6i,l**).

In contrast to that seen in axon nodes, ganglion nodes consistently shortened (**Figure 6g-l**). This pattern is clearly defined in the middle turn across the five critical time periods we analyzed, ranging from P7 to 1M [F(4,29)=19.88, p<0.0001]. Differences in temporal profiles of axonal and ganglion lengths can be explained, in part, by when myelination completed, as shown in **Figure 1**. This explanation is especially true for the ganglion nodes, which have one flanking Schwann-cell paranode and one flanking satellite-cell paranode. Myelination of the somata by satellite cells occurs later than that of the axonal afferents by Schwann cells. We (**Supplementary Figure 1**) and others (Petitpré et al., 2018; Shrestha et al., 2018; Sun et al., 2018) found that differentiation of type I SGNs into three molecularly distinct subtypes occurs between P14 and P21. This differentiation may involve molecular and structural changes that further refine nodal microdomains, leading to differential changes in the length of axonal and ganglion nodes. In agreement with the ultrastructural observations described in **Figure 5**, which suggests that the length of the ganglion node is longer than that of the axon node, our node length measurements show that ganglion nodes are indeed longer than axonal nodes at P7, P10, P14, P21, and 1M and for all cochlear turns measured (**Table 2**).

### The lengths of axonal and ganglion nodes are associated with different aspects of the AN function

During nerve development, adjustments (or refinements) of either myelin thickness or internode length regulate the conduction speed of myelinated axons (Fields, 2008; Kimura & Itami, 2009). This process is involved in synchronization of action potentials (Lang & Rosenbluth, 2003). We hypothesized that changes in axonal and ganglion node lengths are critical for AN development, contributing to the improvement of AN function and hearing onset. To test this hypothesis, we first performed a comprehensive analysis of AN function at P14 and P21 using multi-metric measurements of ABR wave I, which is generated by the AN (McClaskey et al., 2020). This multi-metric approach, which was first developed for human studies (Harris et al., 2018), provides a new tool to assess changes in different aspects of peripheral auditory function *in vivo* with an emphasis on suprathreshold hearing and neural synchrony. The differences in myelin and nodal structures between SGN cell bodies and axons suggests potential functional differences. Comprehensive measurements of AN responses at both threshold and suprathreshold levels are critical to distinguish these differences. Using this novel method, we examined suprathreshold AN function including: 1) estimates of neural synchrony via phase-locking value (PLV), and 2) ABR peak amplitude and peak latency and the change in these metrics with increasing level (**Figure 7; Table 3**). We performed the analyses at P14 and P21, critical time points of myelination and ABR threshold maturation and hearing onset (**Figure 1q**). For the ABR recording, an 11.3 kHz stimulus was selected because we observed well-defined patterns in the changes of the nodal length in the middle turn for both the axonal (**Figure 6 b,e)** and ganglion (**Figure h,k**) nodes and because animals across the different time periods demonstrated their best thresholds at this frequency. After the recordings for P14 and P21 mouse groups were completed, cochleae were collected to be used for node length measurements for correlative structure-function analyses.

**Table 3.** Planned Person product-moment correlation analyses

|  | r | p | 95% Confidence Interval |  |
| --- | --- | --- | --- | --- |
|  |  |  | Lower | Upper |
| Axonal Node Length |  |  |  |  |
| Peak Latency | -0.495 | 0.030 | -0.871 | -0.038 * |
| Mean PLV | 0.545 | 0.018 | 0.120 | 0.880 * |
| Amplitude Intensity Slope | 0.370 | 0.088 | -0.116 | 0.700 |
| Ganglion Node Length |  |  |  |  |
| Peak Latency | 0.403 | 0.068 | -0.017 | 0.745 |
| Mean PLV | -0.364 | 0.091 | -0.788 | 0.213 |
| Amplitude Intensity Slope | -0.526 | 0.022 | -0.855 | 0.217 † |

Note: n = 15. All tests were 1-tailed. 95% confidence intervals were computed from a bootstrap protocol that resampled the data 1000 times. \*p<0.05 and the 95% confidence interval does not cross 0. †Either p<0.05 or the 95% confidence interval does not cross 0.

**Figure 7.**
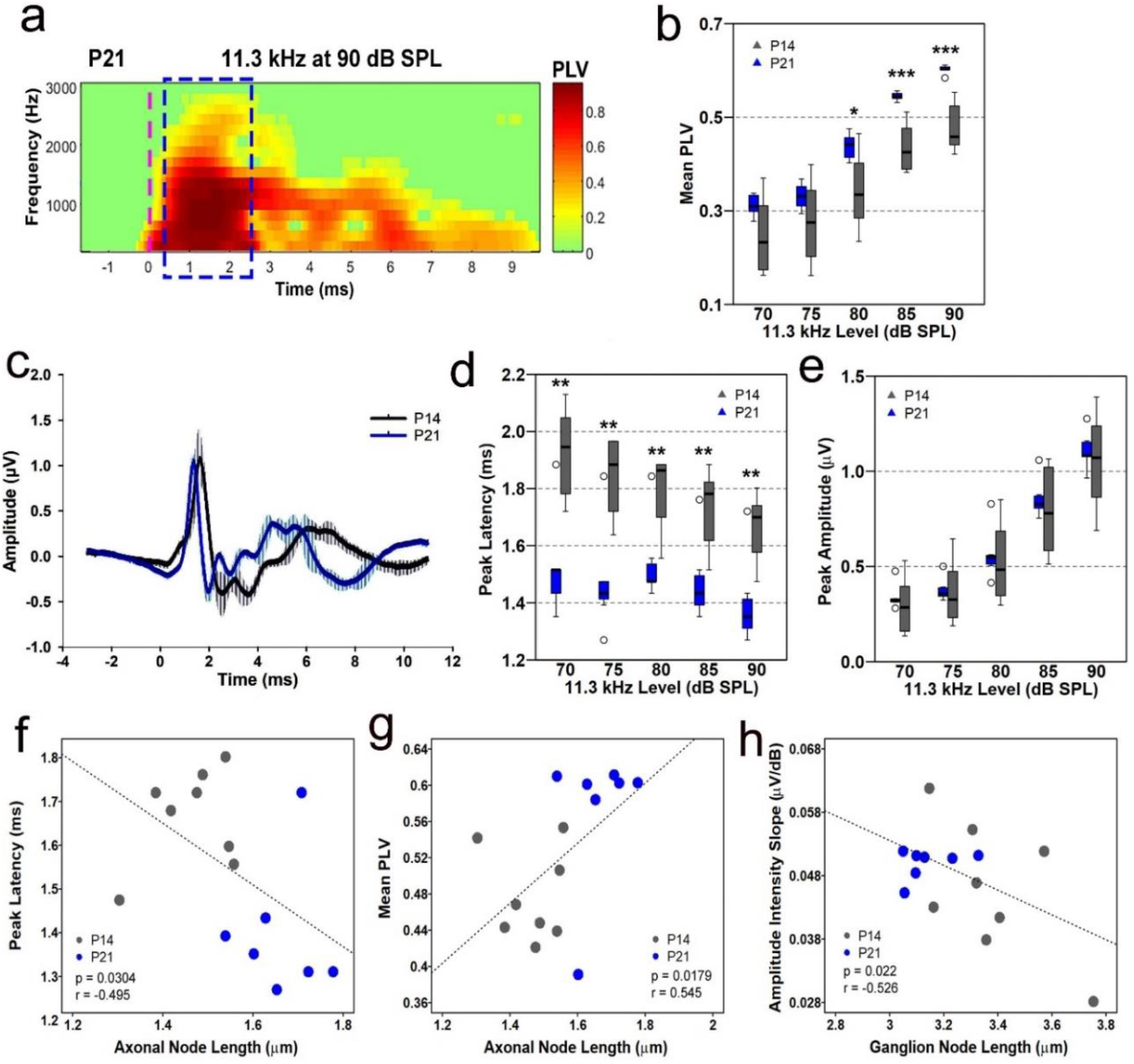
Improvement of AN function evaluated by multiple metrics of ABR and regression analyses between nodal lengths and ABR metrics in the postnatal developing mice. **(a)** A representative example of a time-frequency heatmap of mean phase-locking value (PLV) in a mouse at P21. Dashed blue outline indicates the 2-ms window enclosing wave I where data was collected. The greener shade is closer to baseline, whereas redder shade indicates greater phase-locking. Dashed vertical magenta line indicates stimulus onset. (**b**) Measurements of AN wave I mean PLV between P14 and P21. Synchrony of AN firing of P21 mice is significantly better than in P14 mice at 80-90 dB SPL. **(c)** Group-averaged waveforms of the ABR from postnatal mice at P14 (gray) and P21 (blue) to an 11.3 kHz tone pip at 90 dB SPL; n = 4 mice/group. (**d,e**) Measurements of wave I peak latency (**d**) and amplitude (**e**) between P14 and P21. Peak latencies at P21 are shorter than that of P14 across all levels shown. Peak amplitudes are slightly higher in P21 animals but are not significantly different compared to P14 mice. Two-tailed, Mann-Whitney tests were performed per intensity level between P14 and P21 on **b,d,e**; n(P14) = 8 mice, n(P21) = 7 mice. Outliers are indicated by open circles. * = p<0.05, ** = p<0.01, *** = p<0.001. (**f-h**) Regression analysis across responses at 90 dB SPL from mice at P14 and P21 between the length of axonal nodes and peak latency (**f**) and mean PLV (**g**), and between the length of ganglion node and amplitude intensity slope (**h**). Results of the regression analysis are included in **Table 3**.

Before the correlative analyses, group averaged responses for wave I mean PLV, peak latency, and peak amplitude were compared at each level tested using two-tailed, Mann-WhitneyU testing (**Supplementary Table 4**). Myelination and nodal length development were hypothesized to coincide with faster conduction velocity and greater neural synchrony (shorter latencies and increased PLV, respectively). As predicted, peak latencies of P21 mice are significantly shorter across all intensity levels compared to that of P14 mice (median diff. 90dB = 0.35 ms; p = 0.006) (**Figure 7c,d**). Previous studies found that AN latencies were shorter after hearing onset in BALB/c mice (Song et al., 2006), but synchrony of wave I during the postnatal period of hearing maturation was not determined. As shown in **Figure 7a**, PLV is a unitless measure with a value between 0 and 1, and it represents the consistency of the phase at a given time point surrounding wave I across trials. A value of 0 means no synchrony and a value of 1 means absolute synchrony. At stimulus levels between 80 and 90 dB SPL, the PLVs at P21 are significantly higher than at P14 (median diff. 80dB = 0.11, p = 0.021; 90dB = 0.14, p = 0.0003), revealing better neural synchrony in the older mice (**Figure 7b).** Although peak amplitudes across stimulus levels were not significantly different between P14 and P21 (p = 0.69-0.78), the median response is consistently higher across levels at P21 compared to P14 (range median diffs = 0.01-0.07) (**Figure 7c,e**). The P21 responses, with regards to PLV, latency, and amplitude, are also less variable compared to responses from the P14 group. Increased variability at P14 relative to P21 may be due to individual differences in the temporal pattern of nodal development.

As shown in **Figure 6**, axonal and ganglion nodes display distinctive length patterns during postnatal development. To address the relationship between nodal lengths and AN function during postnatal development, nodal lengths of the axonal or ganglion nodes measured from the middle turn were tested at P14 and P21 for correlation with three ABR metrics, including peak latency, peak amplitude, and PLV obtained at 11.3kHz at 90 dB SPL. Correlation analysis (Pearson’s) was performed using combined data from mice at P14 and P21 (a total of 15 mice) (**Table 3)**. Our analyses revealed that shorter peak latency was significantly correlated to longer axonal nodes (r = 0.495; p = 0.03), whereas no significant correlation was found between the peak latency and length of the ganglion node (r = 0.403; p = 0.068) (**Figure 7f**; **Supplementary Figure 2**). In addition, the length of the axonal node was strongly correlated with higher PLV, with longer axonal node associated with higher PLV (r = 0.545; p = 0.18) (**Figure 7g**). Taken together, these analyses reveal differences in structural development occurring between P14 and P21 result in longer axonal nodes associated with significant decreases in response latency and stronger synchrony. No significant correlation between PLV and ganglion node length was identified (**Supplementary Figure 2**).

The growth in peak amplitude with stimulus intensity, or the amplitude intensity slope, was evaluated in lieu of peak amplitude as it is less susceptible to factors such as head size that can contribute to differences in amplitude across development. Growth in amplitude with increasing level is thought to represent the recruitment of synchronized activity from additional nerve fibers, including both higher threshold fibers and off frequency fibers that are excited due to the spread of excitation along the basilar membrane. We performed correlation analyses using amplitude intensity slope, and node length. Our data shows that shortening ganglion nodes (r = −0.526, p = 0.022), but not lengthening axonal nodes (r = 0.370, p = 0.088), positively correlate with amplitude growth across increasing intensity levels (**Figure 7h; Supplementary Figure 2**). These data suggest that the ganglion node length, but not the axonal node length, contributes to a steeper amplitude intensity slope. Taken together, these analyses suggest that postnatal development of axon nodes underlies enhanced speed and synchrony of AN conduction, whereas refinement of ganglion nodes is associated with greater recruitment of AN fibers, possibly representing the recruitment of higher-threshold fibers.

### Age-related downregulation of nodal related genes and degeneration of nodal structure in the peripheral AN

Loss/dysfunction of nodal components and demyelination have been associated with neurological symptoms of aging (Hinman et al., 2006) and diseases such as Guillain-Barré syndrome and multiple sclerosis (Susuki, 2013). The relationship between aging and changes in nodal structures of the AN has yet to be elucidated. We tested the hypotheses that nodal structures undergo age-related pathophysiological alterations and that the alterations differ between nodal structure types. Immunostaining analysis of ANs in young adult and aged mice demonstrated that node structural integrity, as identified by with NrCAM (node) and Cntn1 (paranode), deteriorates with age in both axonal and ganglion nodes (**Figure 8a**). In addition, TEM imaging revealed elongation of ganglion nodes in aged mice compared to young-adult mice (**Figure 8b**). This observation was confirmed by node length measurements, which revealed a significantly longer ganglion node in the middle turn of ANs in aged mice compared to young-adult mice (**Figure 8d**). The ganglion node was also significantly narrower in both middle and basal turns (**Figure 8h,i**). However, no significant changes in node length or width occurred in axonal nodes (**Figure 8f,g,j,k**). Results of unpaired, two-tailed t-tests comparing age-related node dimension changes are detailed in **Supplementary Table 6**. RNA-seq profiling of ANs in young-adult and aged mice revealed that 26 of 42 node-related genes (62%) were significantly affected in the aged tissue (**Figure 8l, Supplementary Table 5**). Of these affected genes, 23 were downregulated. These genes included key voltage-gated ion channels and structural molecules of the node and paranode. In support of our previous observations (Xing et al., 2012), mice with nodal pathologies showed a robust loss of suprathreshold function, as shown by reduced peak amplitudes and PLV in aged mice (**Supplementary Figure 3**). These data suggest that nodal pathology contributes to age-related declines in auditory function. Changes in nodal length, particularly in the ganglion node, could emerge as a new predictor of declines in AN function.

**Figure 8.**
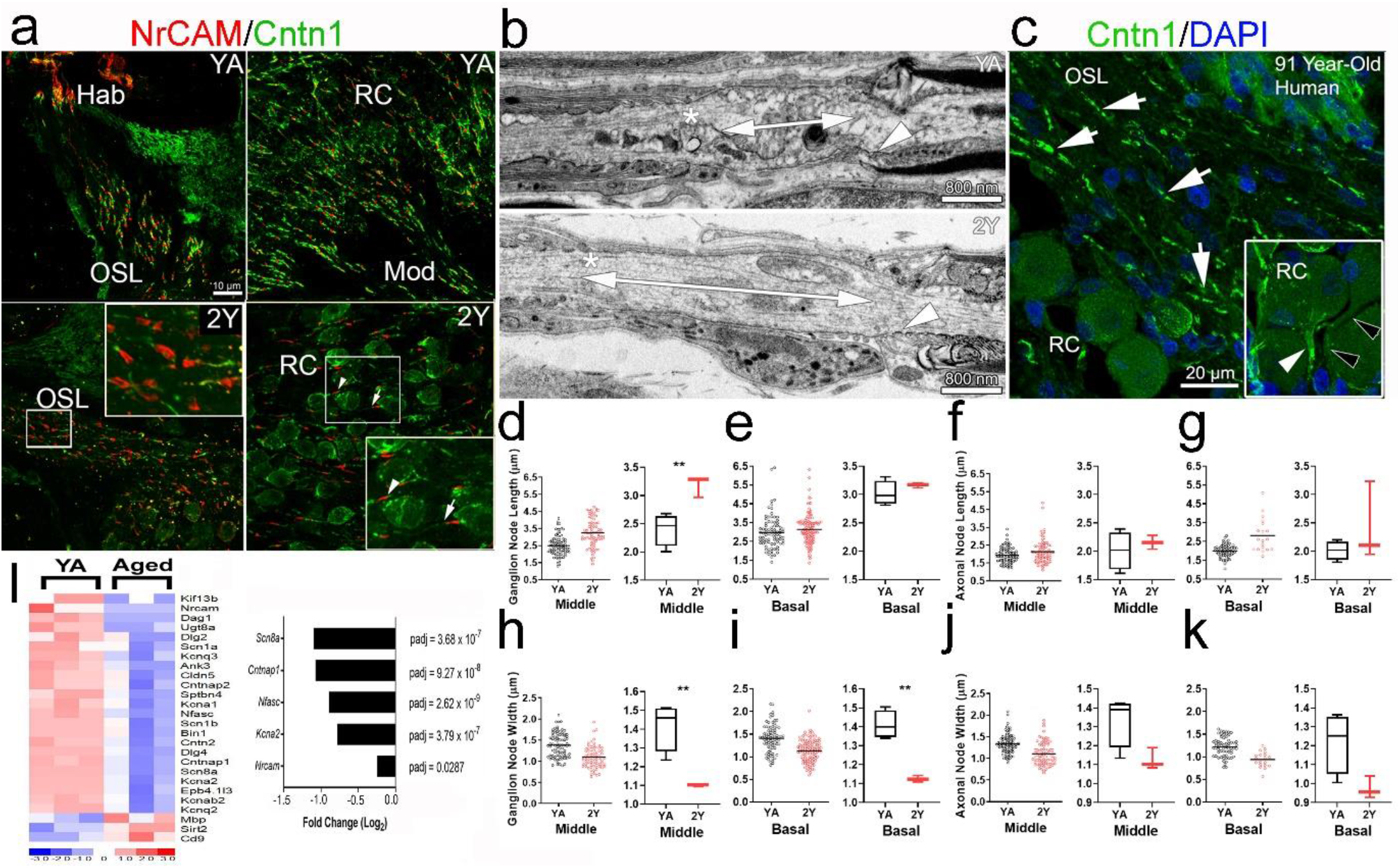
Nodal structures are disrupted in aged ANs. **(a)** Immunostaining for nodal NrCAM and paranodal Cntn1 in ANs of control (YA) and 2-year-old (2Y) mice. Heminode clustering at Hab is sparser in 2Y than in YA (middle right). Axonal node structures are disrupted, with some missing paranodal Cntn1 flanks (bottom-left). The presence of Cntn1 also decreased in some ganglion paranodes (arrow, bottom right) compared to others (arrowhead). (**b**) Transmission electron microscopy imaging revealed an elongated ganglion node in aged (bottom) compared to YA mice (top) as shown by double-ended arrows. Asterisks indicate the satellite cell paranode and arrowheads indicate the Schwann cell paranode. (**c**) Immunostaining for Cntn1 and DAPI of aged human temporal bone showed paranodal flanks (arrows) in human AN. Insert shows Cntn1 in the fiber region preceding the SGN soma (white arrowhead) and another similar fiber region lacking Cntn1 reactivity (black arrowheads). (**d-k**) Measurements of axonal and ganglion node lengths and widths from YA and 2Y ANs. Ganglion node lengths are significantly increased in the middle cochlear turns of 2Y compared to YA and ganglion node widths are significantly decreased in middle and basal turns of 2Y compared to YA. Axonal nodes trend towards elongation and width shrinkage with age. Unpaired, two-tailed t test, ** = p<0.01; nYA = 4 mice, n2Y = 3 mice. Statistical results are detailed in **Supplementary Table 6**. (**l**) Expression profiles of differentially expressed node-related genes show significantly reduced expression with aging compared to YA. The bar graph on the right highlights the negative fold change of several node-related genes of interest; n = 3 mice/group.

## Discussion

Our multidisciplinary approach identified two distinct types of nodes of Ranvier—the axonal node and the ganglion node—in the mouse AN that change across the lifespan, including during myelination and postnatal development, and degenerate during aging. Cellular, molecular, and structure-function correlation evaluations revealed that nodal types are critical for different aspects of auditory nerve function. Here we discuss four key findings pertaining to structural and functional aspects of the node of Ranvier in the AN.

First, the completion of molecular assembly of the nodal structures in the cochlea parallels the onset of hearing function during mouse postnatal development. Clustering of node-related gene expression revealed three stages in node formation: 1) onset of nodal formation at ~P3, 2) complete assembly of structural molecules at ~P14, and 3) further structural and functional maturation between P14 and P21. Transcriptomic findings, together with findings from immunostaining analysis of key structural and functional proteins, nodal NrCAM and paranodal Cntn1, and key voltage-gated sodium channel Na_v_1.6, demonstrate that molecular assembly of the nodes and paranodes was complete by the onset of hearing at approximately P14. Our data also show that NrCAM is present in the basal hook area as early as P3, which supports its importance as an initiator of nodal formation. Previous studies found that NrCAM protein is essential for the initial clustering of voltage-gated sodium channels in the forming nodes and for maintaining and restricting this clustering at the nodes (Amor et al., 2014; Custer et al., 2003; Eshed et al., 2005; Feinberg et al., 2010). Our observations also agree with a recent report describing the maturation of several key voltage-gated ion channels, including Nav1.6, along with the expression of the anchoring protein Ankyrin G and Contactin associated protein 1 in the heminodes and nodes of Ranvier in postnatal rat cochleae between P5 and P7 (Kim & Rutherford, 2016). In addition, our data reveal a temporal-dependence in expression of the axo-glial connector protein Cntn1 in Schwann cell-formed paranodes. These observations demonstrate that molecular assembly of the nodal microdomains begins during early postnatal development and is complete shortly after the onset of hearing function.

Second, we observed two types of nodes of Ranvier in the mouse peripheral AN with distinct features in the nodal and paranodal domains. The structural characteristics of ganglion node, which includes a standard paranodal structure generated by Schwann cells (on the axon side) and a loose paranodal structure provided by satellite cells (on the soma side), was described for the first time in the peripheral auditory system. The unique features of ganglion nodes distinguish them from axonal nodes of the central and peripheral nervous system, such as the nodes of Ranvier of the sciatic nerve, which have provided most of our current knowledge about nodal specializations (see reviews by Peters & Vaughn, 1970; Matthew N. Rasband & Peles, 2016; Salzer, 2015).

Our data also revealed characteristics that distinguish two cochlear glial cells: satellite cells and Schwann cells, which form the paranodal structure and provide myelin sheath to the spiral ganglion cell body and axonal processes, respectively. Myelination of the soma and formation of paranodes by satellite cells are delayed compared to myelination of the axon and formation of paranodes by Schwann cell. Satellite cell myelination of the SGN cell body, although multi-layered, is less dense and especially less compact. This feature is notably visible at the paranodes formed by satellite cells. These observations highlight the role of satellite cells in AN function, which has been understudied. In other nervous systems, satellite cells regulate a wide range of biological processes. For example, satellite cells support neuronal metabolism due to amino acid, nucleobase, and fatty-acid transporters present on their surfaces (Zeisel et al., 2018). Satellite cells are also involved in neuropathic pain (Hanani, 2005; Ohara et al., 2009). In the dorsal root ganglia, myelinating satellite cells expressed fractalkines, which regulate inflammatory responses associated with hypernociception (Souza et al., 2013). In the trigeminal nerve, inhibition of Kir4.1 expression by satellite cells causes dysfunction in potassium buffering, which leads to pain-like behavior (Vit et al., 2008). In the AN, satellite cells but not Schwann cells, also expressed Kir4.1 (Liu et al., 2019), suggesting that satellite cells in the peripheral AN critical regulate AN function. The special feature of the satellite cell-formed paranode in the cochlear nerve (**Figure 4,5,6,7**) provides the structural support for its regulatory function.

Third, our study reveals that from P14 to 1M, while AN function continues to improve after hearing onset, the length of the axonal node in the middle turn continues to increase while the length of the ganglion nodes continues to decrease. Specifically, the length of the axonal node is associated with neural processing speed and neural synchrony, whereas ganglion node development is associated with amplitude growth of the action potential. The middle turn is the most essential part of the sensory epithelium. With the highest nerve fiber density per habenular opening and greatest synapse density per IHC in the middle turn (compared to those in the apex and base), the cochlea has evolved to focus on hearing at the central frequencies (Ehret, 1983; Liberman & Liberman, 2015; Panganiban et al., 2018). Our analyses found significant structure-function correlations between nodal structures and suprathreshold AN function: longer axonal nodes were associated with shorter conduction time and stronger neural synchrony, whereas shorter ganglion nodes were associated with stronger amplitude growth of the AN response. These structure-function analyses reveal distinct roles of cochlear glial cells in regulating AN function. Ganglion nodes and their associated satellite cells relate to amplitude growth, a suprathreshold function of the AN response, whereas axonal nodes and Schwann cells are responsible for AN conduction speed, as measured by peak latency of the ABR wave I response. In the optic nerve and cerebral cortical axon of rats, a computational model showed that the length of the node of Ranvier could determine the velocity of myelinated axon conduction (Arancibia-Cárcamo et al., 2017). A longer axonal node could result in an increased number of voltage-gated sodium channels at the node (if the channel expression level is constant), increasing nerve conduction speed.

To better understand the mechanisms of this striking functional difference between the two types of nodes in the cochlear nerve and their associated glial cells, characterization of the distribution patterns in voltage-gated ion channels in nodal microdomains is needed. Different expression levels of Nav1.6 along the neuron cell body and processes of spiral ganglion was reported in a previous study (Hossain et al., 2005). Using computational modelling, this study concluded that the expression of Nav 1.6 channels at multiple sites along the SGNs contribute to the generation and regeneration of action potentials. The association between ganglion node length and amplitude growth may represent either the recruitment of additional nerve fibers, or the optimization of the generation and regeneration of action potentials with repeated stimulation. Growth in amplitude with increasing sound level is most typically associated with the recruitment of additional nerve fibers, including fibers with higher thresholds (lower spontaneous rate (SR)). The change in ganglion node length and steeper amplitude intensity slope parallels the differentiation of type I AN fibers into the three different subtypes (low-, medium-, and high-SR), and this differentiation was shown to occur approximately between P14 and P21 (Petitpré et al., 2018; Shrestha et al., 2018; Sun et al., 2018) (**Supplementary Figure 1**). Differentiation into multiple type I subtypes may also contribute to the lengthening of axonal nodes post-hearing onset.

Studies of single neuron recording from cats (Liberman & Oliver, 1984), gerbils (Schmiedt, 1989), and mice (Taberner, 2004) show that SGN subtypes have distinct morphologies associated with their functional properties. Mainly, high-SR/low-threshold fibers have larger axon diameters compared to low-SR/high-threshold fibers. Because axon diameter is related to the thickness of myelination, high-SR fibers may be more myelinated than low-SR fibers. Aside from lengthening of the axonal nodes, we also found that the range of axonal node lengths from minimum to maximum is much more variable at P21/1M than at P14. This finding may indicate the presence of the different subtypes, possibly with high-SR axonal nodes on the shorter end and low-SR axonal nodes possibly on the longer end of the length spectrum. Our analysis also found a significant correlation between axonal nodal length and PLV, a measure of synchrony of AN activity. This result suggests further structural maturation of axonal nodes is associated with postanal development of different AN subtypes. Further studies are needed to measure the nodes from the different neuron subtypes to determine if each SGN subtype is associated with a certain axonal node length.

Finally, our data showed an age-dependent downregulation of node-related gene expression and disruption of nodal structures, particularly the ganglion node structures associated with satellite cells, in the mouse cochleae. These effects occurred with declines in neural synchrony and suprathreshold function of the AN response. Based on these observations, degeneration and/or dysregulation of cochlear glial cells, especially satellite cells, may contribute to functional declines in AN associated with aging. Previous studies showed that mice with nodal abnormalities also reduced hearing sensitivity. Noise-exposure caused disruption of myelin and satellite paranodes (Panganiban et al., 2018) and axonal paranodes (Tagoe et al., 2014). Also, transient ablation of glial cells led to disruption of heminodal clustering (Wan & Corfas, 2017). The direct link between abnormal nodal structures, especially that of ganglion nodes, and the declines in multiple components of AN function have yet to be studied. Here, our data indicate that dysregulation of the glial cells and associated degeneration of the ganglion node structure are an important and new mechanism of AN dysfunction in age-related hearing loss.

## Materials and Methods

### Animals

All aspects of animal research were conducted in accordance with the guidelines of the Institutional Animal Care and Use Committee of the Medical University of South Carolina (MUSC). CBA/CaJ mice, originally purchased from the Jackson Laboratory (Bar Harbor, ME), were bred in a low-noise environment at the Animal Research Facility at MUSC. The CBA/CaJ mouse strain is commonly used for normal hearing studies due to their lack of genetic mutations related to hearing and their ability to age without progressive hearing loss (Ohlemiller et al., 2010). Postnatal male and female mice aged P0, P 3, P 7, P 10, P 14, P 21, and 1M were used for developmental studies. Young adult (1.5-3 months) and aged (2-2.5 years) animals were used in the age-related hearing loss study. The numbers of animals per experimental group are reported in the figure legends. All mice received food and water *ad libitum* and were maintained on a 12-hour light/dark cycle. Mice with signs of external ear canal and middle ear obstruction or infection were excluded.

### Physiological procedures

For measurements of auditory function, animals were anesthetized via an intraperitoneal injection of a cocktail containing 20 mg/kg xylazine and 100 mg/kg ketamine. Auditory tests were performed in a sound-isolation booth. Equipment for auditory brainstem measurements was professionally calibrated before use with TDT RPvdsEx software (Tucker Davis Technologies, Gainsville, FL) and a model 378C01 ICP microphone system provided by PCB Piezotronics, Inc (Depew, NY). In a closed-field setup, sound stimuli were delivered into the ear canal via a 3–5-mm diameter tube.

Trial-averaged ABRs were performed using Tucker Davis Technologies equipment System III and processed with their SigGen software package (Version 4.4.1) as previously described (Panganiban et al., 2018). Stimuli were evoked at pure-tone frequencies of 4, 5.6, 11.3, 16, 22.6, 32, 40, and 45.2 kHz. The stimuli consisted of 1.1-ms tone pips with cos^2^ rise/fall times of 0.55 ms and were delivered 31 times/s with a period of 32.26 ms. Responses were collected from 90 to 10 dB SPL sound intensity levels, with each succeeding level being reduced by a 5 dB step. The average ABR waveform for each level tested consisted of 256 trials. Results for each mouse were analyzed for wave I response threshold. Thresholds were then averaged at each frequency and the mean ± standard error of the mean were calculated and plotted using Origin 6.0 software (OriginLab Corporation, Northampton, MA).

### Single-trial recording for ABR metrics analysis

The procedure for continuous single-trial recording was modified from our recent report (McClaskey et al., 2020). For the mice at P14 and P21 used for structure-function correlation analyses, recorded ABRs were elicited only at the 11.3kHz frequency. Stimuli were 1.1 ms in duration with 0.55-ms cosine-squared rise-fall times and were presented at a rate of 21 times/s. At least 500 tone pips of each sound level were presented. Continuous ABR responses were stored and processed offline in MATLAB (MathWorks, Inc., Natick, MA) using the EEGlab toolbox (Delorme & Makeig, 2004) and the ERPlab extension (Lopez-Calderon & Luck, 2014). Continuous data were bandpass filtered between 100 Hz and 3000 Hz using an 8^th^ order Butterworth filter. Data were then epoched from −3 to +11 ms relative to stimulus onset and baseline corrected using the baseline subtraction method with a pre-stimulus window of −3 to −1 ms. Epochs with voltages exceeding +/− 5 μV were rejected and remaining epochs were visually inspected for excessive movement or other artifacts. Artifact-free trials were saved in single-trial form in EEGlab and as a trial-averaged ABR waveforms in ERPlab. Wave I was visually identified as the first major positive inflection following stimulus onset. Peak amplitude and peak latency were calculated from the trial-averaged ABR waveform and PLV was calculated from the single-trial-level data. All measurements were made offline in MATLAB using custom EEGlab and ERPlab functions. Suprathreshold metrics were quantified from responses at 65 dB SPL and above.

Wave I peak amplitude was calculated as the absolute voltage of the wave I peak in microvolts and peak latency was calculated as the latency in ms of the wave I peak. Fractional peak latency (Luck, 2014) is the latency of the point where waveform amplitude has reached 10% of the distance between the wave I start (defined as the local minimum preceding wave I) and wave I peak. The 10% point was used to eliminate the influence of random fluctuations in the start of the waveform (Harris et al., 2018). PLV, also known as inter-trial coherence (ITC), is the length of the vector that is formed by averaging the complex phase angles of each trial at each frequency, and was obtained via time-frequency decomposition of the single-trial-level data. Time-frequency decomposition was performed with Hanning FFT tapers via EEGlab’s newtimef() function, using 16 linearly spaced frequencies from 190 to 3150 Hz. PLV at each timepoint and each frequency was then calculated by taking the absolute value of the complex inter-trial coherence output of the newtimef() function. Mean PLV between 250 and 2500 Hz in the 2-ms window surrounding the wave I peak was then taken as the metric of PLV. A sample PLV time-frequency decomposition is shown in **Figure 7a**.

### Cochlear nerve tissue collection and total RNA isolation

Cochleae were collected at their designated end-points. Microdissections were performed to isolate the AN from the rest of the cochlear structures, taking care to preserve peripheral fibers. Two cochleae from each mouse were pooled for individual samples. Total RNA was purified from each sample using the miRNeasy Mini Kit (Qiagen Inc, Germantown, MD) per the manufacturer’s instructions. The Quality of each total RNA isolation was assessed using the Agilent 2100 Bioanalyzer (Agilent Technologies, Santa Clara, CA). Low-quality samples showing degradation or contamination were excluded.

### RNA sequencing

Total RNA (50 ng) with RNA integrity numbers of 8.9-10 were used for RNA-seq. Samples from P3, 7, 14, and 21 mice were analyzed with each group consisting of three biological replicates. Library preparation and paired-end next-generation sequencing using the Illumina HiSeq2500 (San Diego, CA) was performed by the MUSC Genomics Shared Resource. Resulting FASTq data was processed with Partek Flow™ software. This included alignment with TopHat2, quantification to annotation model (Partek E/M), and normalization and comparison by DeSeq2 (Love et al., 2014). RNAseq datasets were deposited in the NCBI Gene Expression Omnibus (accession numbers GSE141865 and GSE133823). A list of node-related genes was compiled using from manual literature search (Boyle et al., 2001; Custer et al., 2003) and query of amiGO (Carbon et al., 2009) using the terms “node of Ranvier,” “paranode region of axon,” “paranodal junction assembly,” “juxtaparanode,” and “internode.”

### Quantitative PCR

Two-step reverse transcription qPCR (RT-qPCR) was performed by first converting total RNA into cDNA via QuantiTect Reverse Transcription Kit (Qiagen). Resulting cDNA was queried for expression of genes of interest and the 18s reference gene via qPCR using TaqMan probes (Applied Biosystems) and a Lightcycler 480 (Roche Diagnostics). Each experiment included a “no reverse transcription” control; reactions were performed in technical triplicate with 40 cycles of 2-step cycling. Amplification efficiency was calculated by standard curve method using 10-fold serial dilutions of cDNA for each target gene; efficiency was determined by the equation *E* = 10 (−1/S), with S as the slope. Relative expression was calculated using the ∆∆CT method adjusted for calculated efficiencies; measures were normalized based on 18s amount (Livak & Schmittgen, 2001).

### Immunohistochemistry

Immunohistochemical procedures were modified from previous studies (Panganiban et al., 2018). After end-point physiological recordings, mouse cochleae were collected and immediately fixed with 4% paraformaldehyde solution in 1x phosphate-buffered saline for 2 hours at room temperature and decalcified with 0.12M ethylenediamine tetraacetic acid (EDTA) at room temperature for 1-2 days. To prepare for mouse cochlear section, the cochleae were embedded in Tissue-Tek OCT compound and sectioned at a thickness of approximately 10 μm. For whole-mount preparations of mouse cochlear tissues, the ANs and attached organs of Corti were isolated from the rest of the cochlear structures. Sections of human temporal bones were obtained from those we collected in previous studies (Hao et al., 2014). Procedures for human cochlear tissue were reported in detail previously (Hao et al., 2014).

The primary and secondary antibodies used for immunohistochemistry are listed in **Supplementary Table 7**. Staining was performed by either indirect method using biotinylated secondaries conjugated with fluorescent avidin (Vector Labs, Burlingame, CA) or direct method using Alexa Fluor Dyes (ThermoFisher Scientific, Waltham, MA). Nuclei were counterstained using 4′,6-diamidino-2-phenylindole (DAPI). Slice and confocal image stacks were collected using a ZEISS LSM 880 NLO with a ZEN acquisition software (ZEISS United States, Thornwood, NY). Image stacks were taken at 0.75μm intervals with image sizes of 134.95 μm (x) x 134.95 μm (y). Images were processed using ZEN 2012 Blue Edition (Carl ZEISS Microscopy GmbH) and Adobe Photoshop CC (Adobe Systems Incorporated, San Jose, CA).

### Identification of nodal structures and measurement of node lengths

Nodal microdomains were marked via immunostaining of either NrCAM or Na_v_1.6. Paranodal microdomains were marked via immunostaining of Cntn1. Node measurements were acquired using the measurement tool in ZEN. Confocal image stacks taken at 0.75μm intervals from 10μm cochlear frozen sections that were processed into a single-layer maximum intensity projection. Length was taken as the distance between the extreme ends of the immunologically marked nodes (see **Supplementary Figure 4**). Fully-intact nodes in the parallel plane were measured to limit possible inaccuracies. Node parallelness and wholeness were checked using (1) the presence of flanking paranodes, (2) 3D visualization of the tissue section using ZEN, and (3) the range-indicator tool on ZEN.

### Transmission electron microscopy

Samples were prepared for TEM using procedures modified from previous publications (Lang et al., 2015). Briefly, deeply anesthetized mice were cardiac perfused with a mixture of 10 mL saline and 0.1% sodium nitrite solution, followed by 15 mL of a fixative solution containing 4% paraformaldehyde and 2% glutaraldehyde in 0.1 M phosphate buffer, pH 7.4). The same fixative solution was used to perfuse the excised cochleae through the round window and for further immersion overnight at 4°C. Cochleae were then decalcified using 0.12M EDTA solution at room temperature for 2–3 days with a magnetic stirrer. Then, cochleae were fixed using a solution containing 1% osmium tetroxide and 1.5% ferrocyanide for 2 hours in the dark. They were then dehydrated and embedded in Epon LX 112 resin. Semi-thin sections for pre-TEM observation of AN orientation were cut at 1-μm thickness and stained with toluidine blue. Once a coronal plane for a given cochlear turn was seen, ultra-thin sections at 70-nm thickness were cut and stained with uranyl acetate and lead citrate. These ultra-thin sections were examined using a JEOL JEM-1010 transmission electron microscope (JEOL USA, Inc., Peabody, MA).

### Statistical analyses

Sample numbers (n) are indicated in each figure legend. The n for physiological measurements (e.g., amplitude and latency) and RT-qPCR studies were based on prior studies (Lang et al., 2015; Xing et al., 2012). For RNA-seq, prior analyses of mouse AN demonstrated that a sample size of n = 3 was sufficient for robust detection of differentially expressed genes (Lang et al., 2015). Data distribution was tested using either Shapiro-Wilk or Kolmogorov-Smirnov Tests for normality. The appropriate parametric or non-parametric tests were then used. Statistical software and packages used in this project include DeSeq2 (Love et al., 2014), Microsoft Excel, R version 3.5.2 (R Foundation for Statistical Computing, 2019) or GraphPad Prism 8 (GraphPad Software, Inc., La Jolla, CA). For t tests and correlation analyses, a *p*-value of ≤ 0.05 was considered significant. For one-way ANOVAs followed by Benjamini-Hochberg-correction, a q-value of ≤ 0.05 was considered significant. For differential expression analyses of the RNA-seq datasets, a p-adjusted value of ≤ 0.05 was considered significant.

## Acknowledgements

This work was supported by National Institutes of Health Grants R01DC012058 (H.L.), P50DC000422 (H.L., K.C.H.), SFARI Pilot Award (#649452) (H.L), R25 GM072643 (C.H.P., K.V.N.), T32 014435 (C.M.M), P30GM103342 (J.L.B.), P20GM103499 (J.L.B.), P30 CA138313, and S10 OD018113 from Cell & Molecular Imaging Shared Resource and Hollings Cancer Center, C06 RR014516 from the Extramural Research Facilities Program of the National Center for Research Resources, and the Medical University of South Carolina Office of the Vice President for Research. We thank Juhong Zhu and Nancy Smythe for their excellent technical assistance, for Richard Schmiedt for his help with the system setup of single-trial ABR recording, and Jayne Ahlstrom and Crystal Herron for their comments and editing of the manuscript.

