## Supplemental Table 6 for "Two distinct types of nodes of Ranvier support auditory nerve function in the mouse cochlea"

**Supplementary Table 6. Unpaired, two-tailed t test for age-related node length and width changes**

| Axonal Node Length | | | | | |
| --- | --- | --- | --- | --- | --- |
| Turn | Mean (YA) | Mean (2Y) | Mean Diff ± SEM | 95% CI | *p* value |
| Middle | 2.012 | 2.156 | 0.144 ± 0.206 | -0.384 to 0.672 | 0.515 |
| Basal | 2.012 | 2.428 | 0.415 ± 0.353 | -0.492 to 1.322 | 0.292 |
| Axonal Node Width | | | | | |
| Turn | Mean (YA) | Mean (2Y) | Mean Diff ± SEM | 95% CI | *p* value |
| Middle | 1.335 | 1.125 | -0.210 ± 0.084 | -0.428 to 0.008 | 0.056 |
| Basal | 1.219 | 0.974 | -0.245 ± 0.100 | -0.500 to 0.011 | 0.057 |
| Ganglion Node Length | | | | | |
| Turn | Mean (YA) | Mean (2Y) | Mean Diff ± SEM | 95% CI | *p* value |
| Middle | 2.403 | 3.188 | 0.785 ± 0.193 | 0.287 to 1.282 | 0.010 |
| Basal | 3.022 | 3.165 | 0.143 ± 0.128 | -0.186 to 0.472 | 0.315 |
| Ganglion Node Width | | | | | |
| Turn | Mean (YA) | Mean (2Y) | Mean Diff ± SEM | 95% CI | *p* value |
| Middle | 1.417 | 1.099 | -0.318 ± 0.076 | -0.512 to -0.124 | 0.008 |
| Basal | 1.411 | 1.125 | -0.285 ± 0.045 | -0.401 to -0.170 | 0.001 |
