## Supplemental Table 7 for "Two distinct types of nodes of Ranvier support auditory nerve function in the mouse cochlea"

**Supplementary Table 7. Antibodies used in the study**

| **Primary Antibody** | **Host** | **Company** | **Catalog No.** | **Concentration** |
| --- | --- | --- | --- | --- |
| Anti-Cntn1 | Polyclonal goat IgG | R&D Systems (Minneapolis, MN) | AF904 | 1:200 |
| Anti-NF200 | Monoclonal mouse IgG1 | Sigma-Aldrich (St. Louis, MO) | N0142 | 1:200 |
| Anti-NrCAM | Polyclonal rabbit IgG | Abcam (Cambridge, MA) | ab24344 | 1:200 |
| Anti-Calb2 (Calretenin) | Goat | Swant (Switzerland) | CG1 | 1:1000 |
| Anti-Nav1.6 | Polyclonal rabbit | Alomone Labs (Israel) | ASC-009 | 1:100 |
| Anti-Calb1 | Polyclonal rabbit | Cell Signaling (Danvers, MA) | 13176S | 1:100 |
| **Secondary Antibody** | **Host** | **Company** | **Catalog No.** | **Concentration** |
| Anti-Goat Alexa Fluor® 488 | Polyclonal donkey IgG (H+L) | Thermo Fisher Scientific (Waltham, MA) | A-11055 | 1:500 |
| Anti-Rabbit Alexa Fluor® 568 | Polyclonal donkey IgG (H+L) | Thermo Fisher Scientific (Waltham, MA) | A-11011 | 1:500 |
| Biotinylated Anti-Goat IgG | Horse | Vector Laboratories (Burlingame, CA) | BA-9500 | 1:100 |
| Biotinylated Anti-Rabbit IgG | Horse | Vector Laboratories (Burlingame, CA) | BA-1100 | 1:100 |
| Biotinylated Anti-Mouse IgG | Goat | Vector Laboratories (Burlingame, CA) | BA-9200 | 1:100 |
| **Anti-Biotin Dyes** | **Host** | **Company** | **Catalog No.** | **Concentration** |
| Fluorescein Avidin DCS | N/A | Vector Laboratories (Burlingame, CA) | A-2011 | 1:100 |
| Texas Red® Avidin D | N/A | Vector Laboratories (Burlingame, CA) | A-2006 | 1:100 |
