## Supplemental Table 4 for "Two distinct types of nodes of Ranvier support auditory nerve function in the mouse cochlea"

**Supplementary Table 4. Two-tailed, Mann-Whitney tests of supra-threshold differences between P14 and P21**

| Mean PLV | | | |
| --- | --- | --- | --- |
| Level (dB) | Median (P14) | Median (P21) | *p* value |
| 90 | 0.458 | 0.603 | 0.0003 |
| 85 | 0.425 | 0.544 | 0.0003 |
| 80 | 0.335 | 0.442 | 0.021 |
| 75 | 0.275 | 0.332 | 0.281 |
| 70 | 0.233 | 0.310 | 0.094 |
| Peak Latency | | | |
| Level (dB) | Median (P14) | Median (P21) | *p* value |
| 90 | 1.700 | 1.352 | 0.006 |
| 85 | 1.782 | 1.434 | 0.004 |
| 80 | 1.864 | 1.475 | 0.004 |
| 75 | 1.884 | 1.434 | 0.003 |
| 70 | 1.946 | 1.516 | 0.002 |
| Peak Amplitude | | | |
| Level (dB) | Median (P14) | Median (P21) | *p* value |
| 90 | 1.071 | 1.082 | 0.694 |
| 85 | 0.780 | 0.828 | 0.613 |
| 80 | 0.483 | 0.548 | 0.779 |
| 75 | 0.326 | 0.357 | 0.779 |
| 70 | 0.285 | 0.320 | 0.779 |
