## Supplemental Figures 1-4 for "Two distinct types of nodes of Ranvier support auditory nerve function in the mouse cochlea"

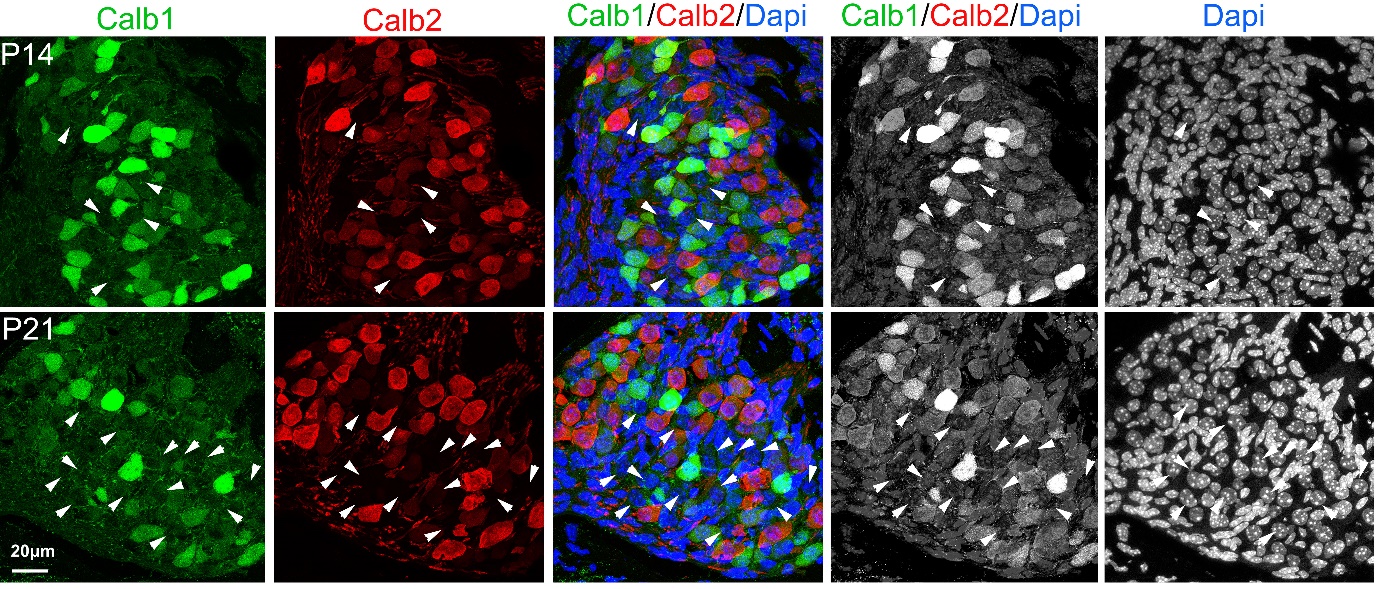
**Supplementary Figure 1**

**Supplementary Figure 1**. Different type I SGN subtype populations in mice at P14 (top panels) versus P21 (bottom panels). High-spontaneous-rate fibers were marked with Calb1, and mid-spontaneous-rate fibers were marked with Calb2. White arrows highlight a subpopulation (low-spontaneous-rate fibers) of SGNs that are Calb1^-^/Calb2^-^.

**Supplementary Figure 2.**

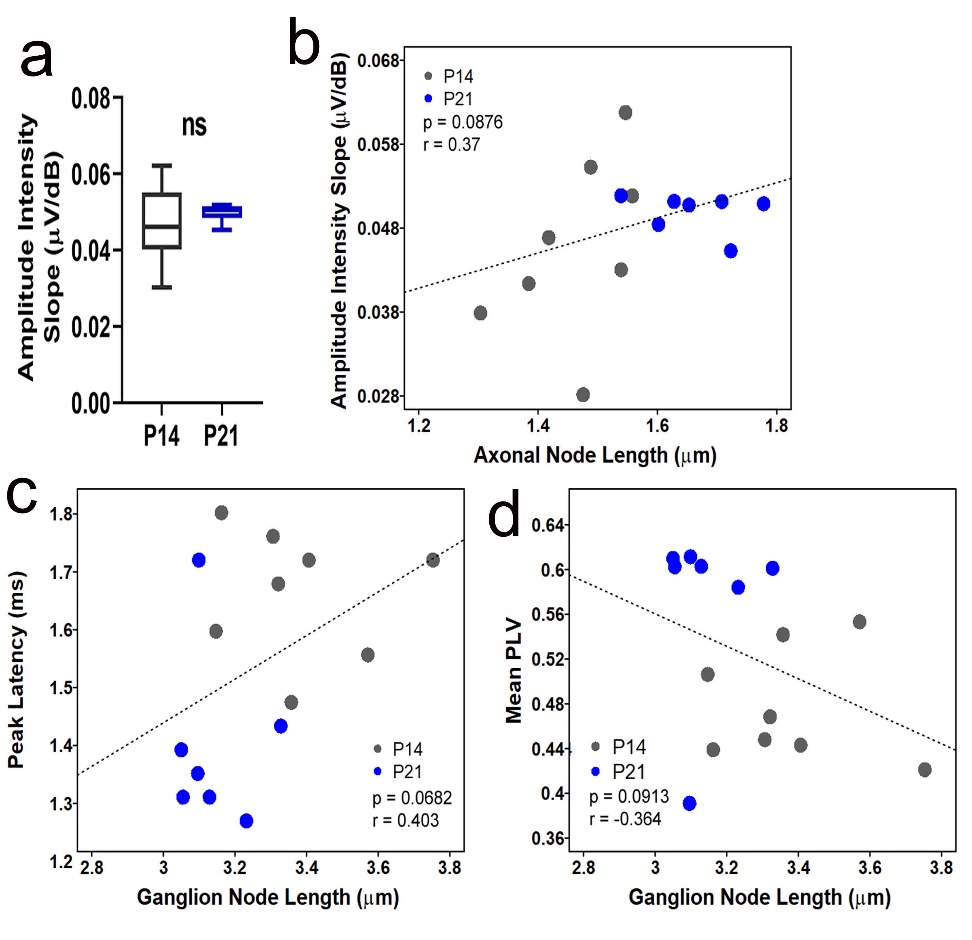

**Supplementary Figure 2.** The slopes of amplitude as a function of stimulus intensity from mice at P14 and P21 and other regression analyses between the nodal lengths and ABR metrics in postnatal developing mice. (**a**) The slope of the amplitudes as a function of stimulus intensity between P14 and P21 groups are not significantly different. n_(P14)_ = 8 mice, n_(P21)_ = 7 mice. (**b**) Correlation analysis across age groups show that axonal node lengths are weakly correlated with amplitude growth, but the association was not significant. Correlation analyses across age groups showed that shorter ganglion node are weakly correlated with reduced peak latencies (**c**) and greater AN synchrony (**d**), but these associations are not significant.

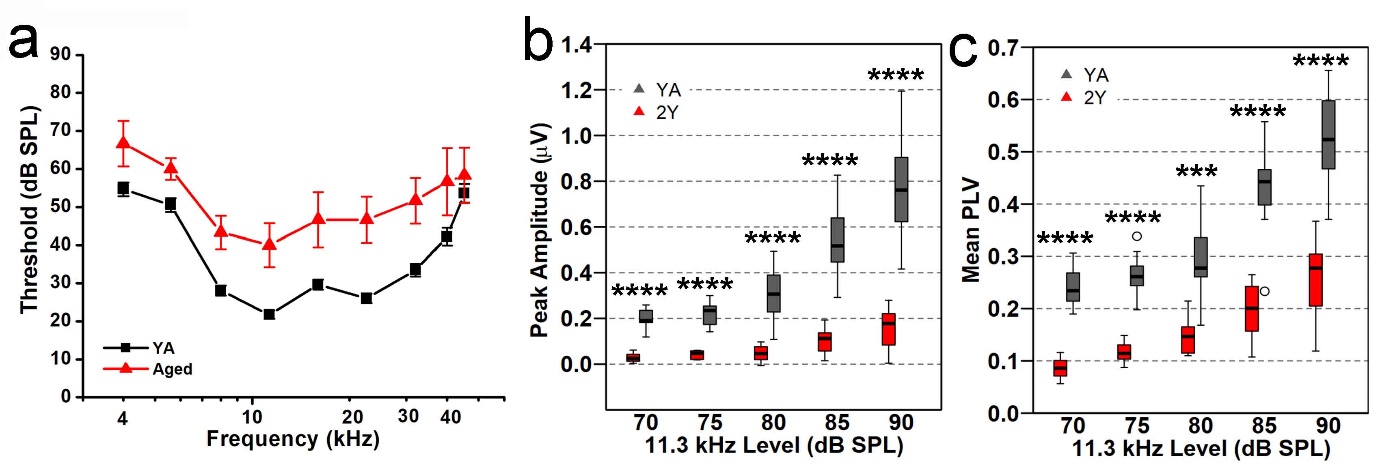
**Supplementary Figure 3.**

**Supplementary Figure 3.** AN responses are smaller in magnitude and dyssynchronous in aged mice**.** (**a**) Audiogram of YA vs 1.5-year-old (1.5Y) aged mice; n_YA_ = 29 mice, n_Aged_ = 3 mice. (**b**) Box plots show greater wave I peak amplitudes from YA compared to 2Y(**c**) Box plots of mean PLV measurements from ABR wave I show that AN responses are more synchronous in YA compared to 2Y mice. Unpaired, two-tailed t-tests *** p < 0.001, **** p < 0.0001; n_YA_ = 10 mice, n_2Y_ = 6–7 mice; open circles indicate outliers.

**Supplementary Figure 4**

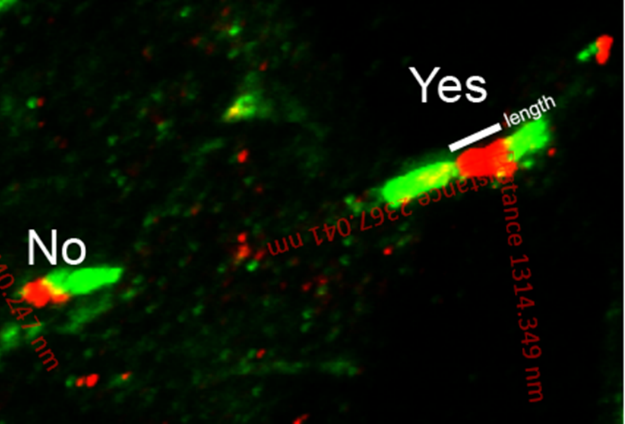

**Supplementary Figure 4.** Node length measurement criteria. This figure shows an example of a parallel axonal node that is good for measuring (indicated by “Yes”) and an axonal node that is not good for measuring (indicated by “No”). The excitable nodal domain is marked by immunostaining for NrCAM (red) and the paranodal flanks are marked by staining for Cntn1 (green).
